# The Essential Facts of Wuhan Novel Coronavirus Outbreak in China and Epitope-based Vaccine Designing against COVID-19

**DOI:** 10.1101/2020.02.05.935072

**Authors:** Bishajit Sarkar, Md. Asad Ullah, Fatema Tuz Johora, Masuma Afrin Taniya, Yusha Araf

## Abstract

Wuhan Novel Coronavirus disease (COVID-19) outbreak has become a global outbreak which has raised the concern of scientific community to design and discover a definitive cure against this deadly virus which has caused deaths of numerous infected people upon infection and spreading. To date, no antiviral therapy or vaccine is available which can effectively combat the infection caused by this virus. This study was conducted to design possible epitope-based subunit vaccines against the SARS-CoV-2 virus using the approaches of reverse vaccinology and immunoinformatics. Upon continual computational experimentation three possible vaccine constructs were designed and one vaccine construct was selected as the best vaccine based on molecular docking study which is supposed to effectively act against SARS-CoV-2. Later, molecular dynamics simulation and in silico codon adaptation experiments were carried out in order to check biological stability and find effective mass production strategy of the selected vaccine. Hopefully, this study will contribute to uphold the present efforts of the researches to secure a definitive treatment against this lethal virus.

## 1. Introduction

### 1.1. Origin of Coronavirus, Their Morphology, Pathology and Others

Coronaviruses (CoVs) are enveloped positive-sense RNA viruses, which is surrounded by crown-shaped, club-like spikes projection on the outer surface [1][2]. Coronaviruses spike protein are glycoprotein that are embedded over the viral envelope. This spike protein attaches to specific cellular receptors and initiates structural changes of spike protein, and causes penetration of cell membranes which results in the release of the viral nucleocapsid into the cell [3]. The viral spike protein includes N-terminal, which is crucial for identification of Coronaviruses [4]. Coronaviruses have a large RNA genome in the size ranging from 26 to 32 kilobases and capable of obtaining distinct ways of replication [5]. Like all other RNA viruses, coronavirus undergoes replication of genome and transcription of mRNAs upon infection. Synthesis of a full-length negative-strand RNA serves as template for full-length genomic RNA [3][6]. Coronaviruses are a large family of viruses belonging to the family Coronaviridae and the order Nidovirales [7]. that are common in many different species of animals, including camels, cattle, cats, and bats [8]. This group of viruses can cause wide varieties of respiratory illness in mammals and birds. In human, it causes common cold, leading to severe illness pneumonia to elderly people and immunocompromised people, like hospital patients [9]. This viral pathogen was responsible for the Middle East Respiratory Syndrome (MERS) and Severe Acute Respiratory (SARS) and 2019 Novel coronavirus (SARS-CoV-2) in China outbreaks [10]. Coronavirus in form of SARS, MERS and Novel coronavirus are lethal leading to large number of deaths. Coronavirus subfamily is classified into four genera, the alpha, beta, gamma and delta coronaviruses. Human Coronavirus (HCoV) infections are caused by alpha- and beta-CoV. CoVs are common human pathogens, and 30% to 60% of the Chinese population is positive for anti-CoV antibodies [11]. These viral infections caused by CoV are generally associated the upper respiratory tract to lower respiratory tract. Immunocompromised people like elderly people and infants are more vulnerable and susceptible to this group of viruses [12].

For many decades HCoVs are well adapted to humans and it had been prevalent among Human races. In 1960s, the first identification of Human coronaviruses (HCoVs) was made in patients with common cold. Following that incident more HCoVs has been detected, that involved the two major outbreak SARS and MERS, two pathogens that, upon infection, can cause fatal respiratory disease in humans [13].

The most common coronaviruses among humans are 229E, NL63, OC43, and HKU1 and some can evolve and cause human diseases, becoming new human coronaviruses. Three recent examples of these are SARS-CoV, SARS-CoV-2 and MERS-CoV [14]. SARS-CoV was the causal agent of the severe acute respiratory syndrome outbreaks in 2002 and 2003 in Guangdong Province, China. 6-8 MERS-CoV was the pathogen responsible for severe respiratory disease outbreaks in 2012 in the Middle East [15].

Chinese scientists were able to sequence the viral genome using next genome sequencing from the collected samples and identified the cause as a SARS-like coronavirus. The genetic sequence is now available for scientist all around the world which would aid faster diagnosis of further cases [2] [16][17]. Based on full genome sequence data on the Global Initiative on Sharing All Influenza Data [GISAID] platform, the genetic sequence of the SARS-CoV-2 ensuring faster development of point-of-care real-time RT-PCR diagnostic tests specific for novel Coronavirus diseases or COVID-19 [18]. Association of genome sequence with other evidences demonstrated that SARS-CoV-2 is 75 to 80% identical to the SARS-CoV and even more closely related to several bat CoVs. SARS-CoV-2 can be cultured in the same cells that are ideal for growing SARS-CoV and MERS-CoV [19]. Another group of scientists have investigated the origin of the SARS-CoV-2, using comprehensive sequence analysis and comparison in conjunction with relative synonymous codon usage (RSCU). The evidence and analysis suggest that the SARS-CoV-2 appears to be a recombinant virus between the bat coronavirus and an origin-unknown coronavirus. the viral spike glycoprotein recombined within themselves, and recognizes cell surface receptor. Scientist are speculating that the SARS-CoV-2 originated form snakes based on Homologous recombination within the spike glycoprotein could be a probable way of transmission in between snake and humans [20].

### 1.2. Severity of China Outbreak of Wuhan Novel Coronavirus (2019-nCoV)

China is in its extremity due to the outbreak of a respiratory virus, called coronavirus and considered as related to SARS virus [21][22]. It is reported to be originated in the animal and seafood market of Wuhan, Hubei province, China – one of the largest cities in China which has 11M residents thus entitled as Wuhan Coronavirus [21] [23]. The disease it caused is called novel Coronavirus disease or COVID-19. World health organization (WHO) is investigating the virus since 31^st^ December, 2019 after being reported [24][25]. SARS-CoV-2 is a deadly virus and Chinese authority reported WHO a pneumonia like case on 31^st^ December, 2019 and the infection has spread destructively since then [26]. First death of a 61 years old man was reported on 11^th^ January by Chinese health authorities in Wuhan, Hubei province, China [25][27][28]. Second death of a 69 years old man was reported on 15^th^ January in the central city of Wuhan, Hubei province, China who had symptoms of pneumonia [29][30]. In Thailand, first emerging international case outside the China was reported on 13^th^ January and the first emerging case outside the Wuhan city, China was reported on 19^th^ January [25][31]. On January 20, it was reported that COVID-19 virus can be transmitted from human to human which cracked the idea that it can only be transmitted through infected animals [32]-[35] and on the same day China Health authority reported the third death [36]. It was recorded that the number of infections rose up to 10 fold between 17^th^-20^th^ January [37]. United States confirmed the first infected case in Washington state on 21^st^ January who had recently been returned from china [38]. As of 23^rd^ January China reported 830 infected cases and the death toll rose up to 25 in China [39][40]. On the same date several confirmed infected cases were reported worldwide, Taiwan (1 case), Macao(2 cases), Hong Kong(2cases),Vietnam (2cases), Thailand (3cases), Japan (2cases), South Korea(2 cases),Singapore (1case), United States (1case) [41]. On 24 January the health minister of France Agnes Buzyn announced three confirmed infected cases, two in Paris and 1 case in Bordeaux. France is the first European country to be announced as infected [42]-[44].

Chinese authority banned public and air transport in Wuhan on 23 January and the residents of the country were kept under quarantine [45]. National Health Commission of China reported on 25^th^ January that as of 24^th^ January the virus took 41 lives and 1300 infected cases were reported worldwide [46]-[48]. As of January 25, China confirmed 2000 infected cases worldwide, among these 1975 cases were reported in China with rising death toll to 56 [49]-[51]. Chinese health authority announced on January 27 that the coronavirus death toll jumped to 81 and 2744 infected cases in china were reported [52]-[54]. German authorities confirmed first infected case of coronavirus and it is reported as second European country to be infected. The health authority said that the patient is in the good condition and is kept under medical observation [55][56]. Health and Human Services Secretary of U.S. said on 28^th^ January that the infected cases in U.S. remained five with no deaths, while France reported forth infected case. In china 106 deaths with more than 4520 infected cases were reported on 28^th^ January [57][58]. Alarming situation due to rising confirmed cases outside China made governments of other countries to start evacuation of their citizens from Wuhan, China and these include U.S., France, Japan, Australia, Germany, New Zealand, France [59]. Approximately, 170 deaths and 7,711 confirmed cases were reported with 1,737 new cases by Chinese National health commission on 29^th^ January [60]. The U.S. government evacuated 240 Americans, mostly diplomats and their family members from the epicenter of the virus outbreak on 29^th^ January by a chartered airplane [61]. Another report stated that on the same day chartered ANA airplane by Japanese government landed in Heneda Airport after evacuating 206 Japanese nationals [62].

The World Health Organization (WHO) had a meeting at the WHO headquarter in Geneva on 30^th^ January and WHO declared the outbreak as Public Health Emergency and it is also said by the committee that the declaration was made giving priority the other countries outside China [63]. On 30^th^ January the U.S. state department advised their citizens not to travel to China as the death toll jumped to 204 with 9,692 infected cases worldwide [64]. In China 213 deaths and globally 9,826 confirmed cases were reported on 31^st^ January by Chinese national health authorities, among which 9,720 cases in China and 106 cases in 19 countries outside China [65][66]. On 1^st^ February the death toll reached to 258 with 11,948 confirmed cases worldwide stated by Hubei Provincial Health Commission [67][68]. On the same day German Ministry of Health said that a military airplane landed at Frankfurt airport after evacuating 128 German citizens from Wuhan who will be quarantined till mid of February and the health minister also stated that Germany military aircraft delivered aid and 1000 protective suits to Chinese authorities requested by China [69].

First known fatality outside China reported of a 44 year old man in Philippines by Philippine health officials on 2^nd^ February as the novel coronavirus outbreak took 362 lives and 17,205 cases were reported according to Chinese and World Health Organization data [70][71].The Nation’s flagship carrier, Qantas, was chartered by Australian government and the airplane landed in Learmonth, Western Australia after evacuating 243 citizens including 89 children on 3^rd^ February from where the citizens would be sent by another flight to Christmas Island to be quarantined [72]. New 1000-bed facility Huoshenshan Hospital in Wuhan is built in only 10 days to mitigate the pressure of patients in other medical hospitals in Wuhan. A crew of more than 7000 workers started working around the clock on 24 January which covers 60000 square meters with special quarantine area and opened its door to receive patients on 3^rd^ February [73][74]. Kerala government announced the third confirmed case in India on 3^rd^ February who is a student [75] just like the 1^st^ case (reported on 30^th^ January in Kerala) and 2^nd^ case (reported on 2^nd^ February in Kerala) according to the Ministry of Health and Family welfare of India [76][77]. A British pharmaceutical company, GlaxoSmithKline (GSK) stated on 3^rd^ February that they would like to collaborate with Coalition for Epidemic Preparedness Innovations (CEPI) that works for developing vaccines. GSK said that they are ready to work with their adjuvant technology to increase immunity to the patients as well as to develop vaccine for coronavirus outbreak Health authorities in China stated on 4^th^ February that as end of 3^rd^ February the coronavirus fatalities jumped to 425 in China and 427 worldwide as the second fatality announced in Hong Kong of a 39 year old man reported on same day in the early morning [75][78]. The authority also said that the confirmed cases reached to 20,438 worldwide [79][80]. COVID-19 claimed second death of a 39-year-old man in Hong Kong, outside mainland China on 4th February, 2020 [81].

On 5th February, 2020 a Japanese cruise ship, docked at Yokohama, Japan called Diamond Princess was quarantined for two weeks by Health officials after the disembarked Hong Kong citizen tested positive of virus which started it’s round trip carrying 3711 individuals on board including 2666 passengers and 1045 crew members [82]. Within 48 hours the number of infected cases on the cruise ship jumped sharply to 61, reported on 7th February, 2020 [83] and another report stated the death of the Chinese doctor, Li Wenliang, who tried to warn about the destructivity of coronavirus on the same date and more than 630 fatalities along with more than 30,000 infected cases were reported worldwide [84][85]. As of 12th February, 2020 the infected cases on the cruise ship increased around 3 fold and the number was 174 and this cruise ship is considered as the large cluster of infected patients outside China by the Health experts and compared with “ Floating petri dish” because it is easier to spread infectious disease in such a confined area[86]. The National Health Commission of China announced 1,843 deaths and 56,249 confirmed cases in Hubei, China on 15th February, 2020 [87]. First European fatality of a 80-year-old-man in France was reported on the same date [88]. 1,933 deaths in Hubei alone and more than 70,000 cases were reported by Health authorities on 18th February, 2020 [89]. China’s National Health Committee reported on 19th February, 2020 that blood plasma can be extracted from the recovered patients to treat serious patients [90]. Japan’s Health Ministry announced two deaths of a 87-year-old-man and 84-year-old-woman on the cruise ship on 20th February, 2020 and on the same date South Korea’s first fatality was reported [91]. The youngest coronavirus infected patient who is under age 10 was reported in Japan on 21th February, 2020 [92]. As of 22th February, 2020, 2,239 death cases and 75,567 infected cases including 346 infected cases in South Korea were reported to the World Health Organization (WHO) [93]. World Health Organization (WHO) welcomed the report of China on 23th February, 2020 saying there was a fall in new death case in mainland China but concerned with infected cases in other counties as the total infected cases in South Korea stood 433 on the same date [94].

To date, there is no effective antiviral therapies that can combat the Coronavirus infections and hence the treatments are only supportive. Use of Interferons in combination with Ribavirin is somewhat effective. However, the effectiveness of combined remedy needs to be further evaluated [95]. This experiment is carried out to design novel epitope-based vaccine against four proteins of SARS-CoV-2 i.e., nucleocapsid phosphoprotein which is responsible for genome packaging and viral assembly [96]; surface glycoprotein that is responsible for membrane fusion event during viral entry [97][98]; ORF3a protein that is responsible for viral replication, characterized virulence, viral spreading and infection [99] and membrane glycoprotein which mediates the interaction of virions with cell receptors [100] using the approaches of reverse vaccinology.

Reverse vaccinology refers to the process of developing vaccines where the novel antigens of a virus or microorganism or organism are detected by analyzing the genomic and genetic information of that particular virus or organism. In reverse vaccinology, the tools of bioinformatics are used for identifying and analyzing these novel antigens. These tools are used to dissect the genome and genetic makeup of a pathogen for developing a potential vaccine. Reverse vaccinology approach of vaccine development also allows the scientists to easily understand the antigenic segments of a virus or pathogen that should be given more emphasis during the vaccine development. This method is a quick, cheap, efficient, easy and cost-effective way to design vaccine. Reverse vaccinology has successfully been used for developing vaccines to fight against many viruses i.e., the Zika virus, Chikungunya virus etc. [101][102].

## 2. Materials and Methods

The current experiment was conducted to develop potential vaccines against the Wuhan novel coronavirus (strain SARS-CoV-2) (Wuhan seafood market pneumonia virus) exploiting the strategies of reverse vaccinology (**Figure 01**).

**Figure 01.**
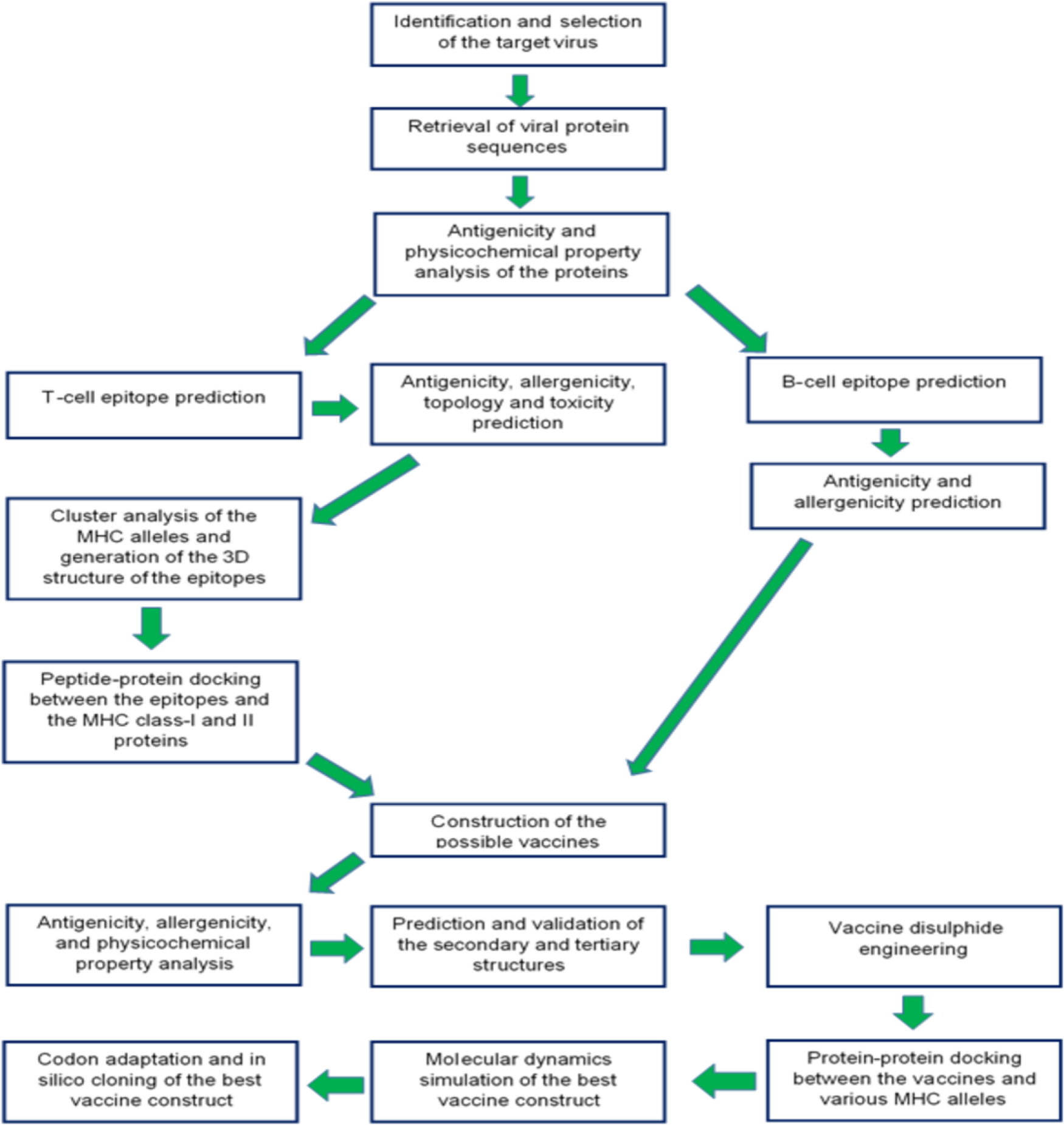
Strategies employed in the overall study

### 2.1. Strain Identification and Selection

The strain of the SARS-CoV-2 was selected by reviewing numerous entries of the online database of National Center for Biotechnology Information or NCBI (https://www.ncbi.nlm.nih.gov/).

### 2.2. Retrieval of the Protein Sequences

Four protein sequences i.e., nucleocapsid phosphoprotein (PubMed accession no: QHD43423.2), membrane glycoprotein (PubMed accession no: QHD43419.1), ORF3a protein (PubMed accession no: QHD43417.1) and surface glycoprotein (PubMed accession no: QHD43416.1) were downloaded from the NCBI (https://www.ncbi.nlm.nih.gov/) database in fasta format.

### 2.3. Antigenicity Prediction and Physicochemical Property Analysis of the Protein Sequences

The antigenicity of protein sequences was determined using an online tool called VaxiJen v2.0 (http://www.ddg-pharmfac.net/vaxijen/VaxiJen/VaxiJen.htm). During the antigenicity prediction, the prediction accuracy parameter threshold was set at 0.4 in the tumor model [103]-[105]. The tumor model was used in prediction because it generated excellent results (when compared to other models) in both the leave-on-out cross-validation (LOO-CV) and external validation in numerous studies. The accuracy, sensitivity, and specificity of a prediction (by the VaxiJen v2.0 server) depends on the prediction accuracy threshold. For this reason, to improve the accuracy of the prediction, the threshold 0.4 was used. The server is a user-friendly web tool for effective prediction of the antigenicity of proteins [106]-[108]. Only the highly antigenic protein sequences were selected for further analysis. Next, the selected antigenic protein sequences were analyzed by ExPASy’s online tool and ProtParam (https://web.expasy.org/protparam/) to determine their physicochemical properties i.e., the number of amino acids, theoretical pI, extinction co-efficient, aliphatic index and grand average of hydropathicity (GRAVY) etc. [109].

### 2.4. Prediction of T-cell and B-cell Epitopes

The online epitope prediction server Immune Epitope Database or IEDB (https://www.iedb.org/) was used for T-cell and B-cell epitope prediction. The database contains a huge collection of experimental data on T-cell epitopes and antibodies [110]. The NetMHCpan EL 4.0 prediction method was used for MHC class-I restricted CD8+ cytotoxic T-lymphocyte (CTL) epitope prediction (for HLA-A*11-01 allele) and the MHC class-II restricted CD4+ helper T-lymphocyte (HTL) epitopes were predicted (for HLA DRB1*04-01 allele), using the Sturniolo prediction method [111]. Ten of the top twenty MHC class-I and MHC class-II epitopes were randomly selected based on their antigenicity scores (AS). For, the B-cell lymphocytic epitopes (BCL), with amino acid number of more than ten, were selected for analysis that were predicted using BepiPred linear epitope prediction method [112] [113].

### 2.5. Transmembrane Topology, Antigenicity Prediction, Allergenicity and Toxicity Prediction of the Selected Epitopes

The epitopes selected in the previous step were then subjected to the transmembrane topology experiment using the transmembrane topology of protein helices determinant, TMHMM v2.0 server (http://www.cbs.dtu.dk/services/TMHMM/) [114]. The antigenicity of the epitopes were determined by the online VaxiJen v2.0 (http://www.ddg-pharmfac.net/vaxijen/VaxiJen/VaxiJen.htm) server. The threshold was kept at 0.4 in the tumor model for improving the accuracy of the prediction [106]-[108].

The allergenicity of the selected epitopes were predicted using two online tools i.e., AllerTOP v2.0 (https://www.ddg-pharmfac.net/AllerTOP/) and AllergenFP v1.0 (http://ddg-pharmfac.net/AllergenFP/). AllerTOP server generates more accurate results than AllergenFP server [115] [116]. For this reason, during prediction, the results predicted by AllerTOP were given much priority. The toxicity prediction of the selected epitopes was carried out using ToxinPred server (http://crdd.osdd.net/raghava/toxinpred/) using SVM (support vector method) based method, keeping all the parameters default. The SVM is a machine learning technique which is used in the server for differentiating the toxic and non-toxic peptides [117]. After the antigenicity, allergenicity and toxicity tests, the epitopes that were found to be antigenic, non-allergenic and non-toxic, were considered as the best predicted epitopes and selected for the later phases of the experiment.

### 2.6. Cluster Analysis of the MHC Alleles

The cluster analysis of the MHC alleles was carried out to identify the alleles of the MHC class-I and class-II molecules with similar binding specificities. The cluster analysis of the MHC alleles were carried out using online tool MHCcluster 2.0 (http://www.cbs.dtu.dk/services/MHCcluster/) [118]. During the analysis, all the parameters were kept default and all the HLA supertype representatives (MHC class-I) as well as the HLA-DR representatives (MHC class-II) were selected. The NetMHCpan-2.8 prediction method was used for analyzing the MHC class-I alleles.

### 2.7. Generation of the 3D Structures of the Selected Epitopes

The 3D structures of the selected epitopes were generated using PEP-FOLD3 (http://bioserv.rpbs.univ-paris-diderot.fr/services/PEP-FOLD3/) online tool. Only the best selected epitopes from previous steps (the epitopes that followed the selection criteria of high antigenicity, non-allergenicity and non-toxicity in the previous steps, were considered best) were taken for 3D structure prediction [119]-[121].

### 2.8. Molecular Docking of the Selected Epitopes

Molecular docking analysis is one of the essential steps in reverse vaccinology to design vaccines. In this step, peptide-protein docking was performed to predict the binding of epitopes with various antibodies or the MHC receptors [122]. The molecular docking study of the epitopes were conducted using PatchDock (https://bioinfo3d.cs.tau.ac.il/PatchDock/php.php) server against HLA-A*11-01 allele (PDB ID: 5WJL) and HLA DRB1*04-01 (PDB ID: 5JLZ). PatchDock tool is a quick, easy, efficient and effective online docking tool which works on some specific algorithms. These algorithms divide the connolly dot surface representations of the compounds into concave, convex and flat patches. Later, possible solutions are predicted from these patches and RMSD (root mean square deviation) scores are applied to the candidate solutions. Based on the RMSD scores, the best candidate solutions are determined by the server. After successful docking by PatchDock, the refinement and re-scoring of the docking results were carried out by the FireDock server (http://bioinfo3d.cs.tau.ac.il/FireDock/php.php). The FireDock server generates global energies for the best solutions and the lowest global energy is always considered as the best docking score [113]-[126]. The best results were visualized using Discovery Studio Visualizer [127].

### 2.9. Vaccine Construction

Three possible vaccines were constructed against the selected Wuhan novel coronavirus. The predicted CTL, HTL and BCL epitopes were joined together by different linkers for vaccine construction. The vaccines were constructed maintaining the sequence: adjuvant, pan HLA DR Epitope (PADRE) sequence, CTL epitopes, HTL epitopes and BCL epitopes. Three different adjuvants i.e., beta defensin, L7/L12 ribosomal protein and HABA protein (*Mycobacterium tuberculosis*, accession number: AGV15514.1), were used and the three vaccines differ from each other by their adjuvant sequence. Beta-defensin adjuvant acts as agonist to stimulate the activation of the toll like receptors (TLRs) 1, 2 and 4. On the other hand, TLR-4 is activated by the L7/L12 ribosomal protein and HABA protein. Studies have also confirmed that, the PADRE sequence stimulates the cytotoxic T-lymphocyte response of the vaccines that contain it. During the vaccine construction, four different linkers: EAAAK, GGGS, GPGPG and KK, were used [128]-[135]. **Figure 02** illustrates how the linkers, epitopes, adjuvants and PADRE sequence were used for vaccine construction, in their appropriate order. The three vaccines differ from each other only in their adjuvant sequences.

**Figure 02.**
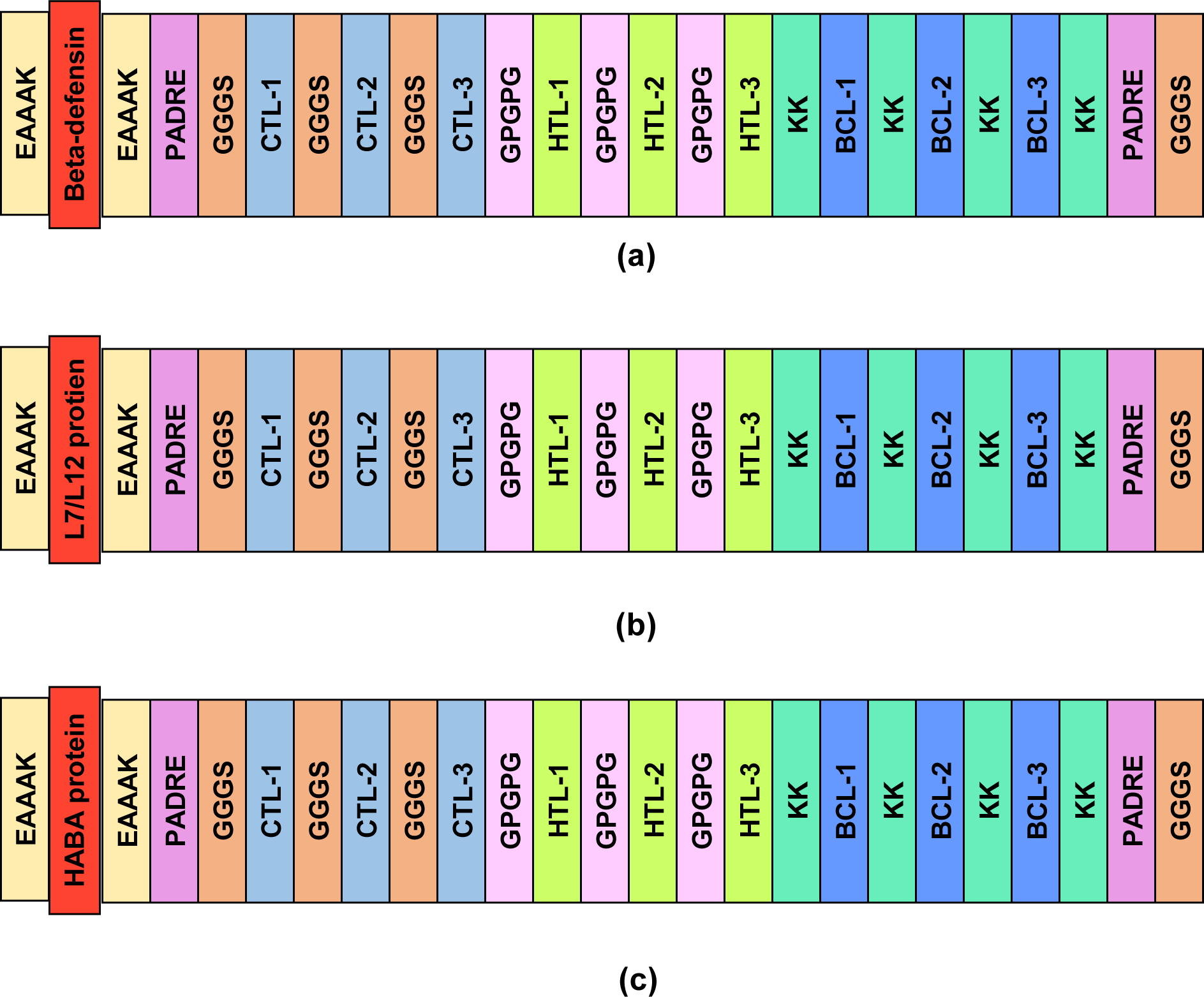
A schematic representation of the three possible vaccine constructs with linkers (EAAAK, GGGS, GPGPG, KK), PADRE sequence, adjuvants (beta-defensin, L7/L12 protein, HABA protein) and epitopes (CTL, HTL, BCL) in sequential and appropriate manner. (a) is the first vaccine constructed using beta-defensin adjuvant, (b) is the second vaccine constructed using L7/L12 adjuvant protein and (c) is the third vaccine constructed using HABA protein as adjuvant. CTL; cytotoxic T lymphocytic epitope, HTL; helper T lymphocytic epitope, BCL; B cell lymphocytic epitope. The three vaccine constructs defer from each other only in their adjuvant sequences.

### 2.10. Antigenicity, Allergenicity and Physicochemical Property Analysis of the Predicted Vaccines

The antigenicity of the constructed vaccines was predicted by the online server VaxiJen v2.0 (http://www.ddg-pharmfac.net/vaxijen/VaxiJen/VaxiJen.htm) using the tumor model where the threshold was set at 0.4. The allergenicity of the predicted vaccines were determined by two online tools for improving the prediction accuracy i.e., AlgPred (http://crdd.osdd.net/raghava/algpred/) and AllerTop v2.0 (https://www.ddg-pharmfac.net/AllerTOP/). MEME/MAST motif prediction was used to predict the allergenicity of the vaccines by AlgPred [136]. Later, ProtParam (https://web.expasy.org/protparam/) was used to predict different physicochemical properties of the predicted vaccines [109].

### 2.11. Secondary and Tertiary Structure Prediction of the Vaccine Constructs

The secondary structure generation of the vaccine constructs were performed using online server, PRISPRED (http://bioinf.cs.ucl.ac.uk/psipred/), which is a simple and easy secondary structure generating tool [137][138]. The secondary structures of the vaccine constructs were predicted using the PRISPRED 4.0 prediction method. Next, another online tool, NetTurnP v1.0 (http://www.cbs.dtu.dk/services/NetTurnP/) was used to predict the β-sheet structure of the predicted vaccines [139]. The 3D structures of the vaccines were then generated using online server RaptorX (http://raptorx.uchicago.edu/) [140]-[142].

### 2.12. 3D Structure Refinement and Validation

Protein structure refinement is an important step in vaccine design because when protein 3D structures are predicted by online tools, they may lack the true, native structure. The refinement of the structures can convert the low resolution predicted model to high resolution predicted model that closely resembles the native structure of the protein. The 3D structures of the constructed vaccines were refined using online protein refinement tool, 3Drefine (http://sysbio.rnet.missouri.edu/3Drefine/). The tool is a quick and efficient 3D structure refining tool (works on i3Drefine protocol) since it can refine a 300 amino acid long protein in just 5 minutes [143] [114]. Next, the validation was conducted by analyzing the Ramachandran plots which were generated using the PROCHECK (https://servicesn.mbi.ucla.edu/PROCHECK/) server [145][146].

### 2.13. Vaccine Protein Disulfide Engineering

Disulfide bonds are important characteristics of protein because they provide stability to the proteins. For this reason, to predict the correct disulfide interactions, the vaccine protein disulfide engineering was carried out with the aid of Disulfide by Design 2 v12.2 (http://cptweb.cpt.wayne.edu/DbD2/) server which predicts the potential sites within a protein structure that have higher possibility of undergoing disulfide bond formation [147]. During disulfide engineering, the intra-chain, inter-chain and Cβ for glycine residue were selected and the χ3 Angle was kept -87° or +97° ± 5 and Cα-Cβ-Sγ Angle was kept 114.6° ±10. The χ3 angle was set at -87° or +97° ± 5 because numerous disulfides were generated by the server, when the default angle (+97° ±30° and -87° ±30°) was used. So, to shorten the amount of disulfides, the χ3 angle was kept low. Furthermore, studies have estimated that Cα-Cβ-Sγ angle reach a peak of near 115° and covers a range from 105° to 125°, in known disulfides. For this reason, the Cα-Cβ-Sγ angle was kept default (114.6° ±10) [148] [149].

### 2.14. Protein-Protein Docking

In protein-protein docking, the constructed SARS-CoV-2 vaccines were analyzed by docking against some MHC alleles as well as the toll like receptors (TLRs). During viral infections, the MHC complex recognize the viral particles as potent antigens. Since the different portions of the MHC molecules are encoded by different MHC alleles, the constructed vaccines should have very good binding affinity towards these segments of the MHC complex [150]. In this experiment, all the vaccines constructs were docked against DRB1*0101 (PDB ID: 2FSE), DRB3*0202 (PDB ID: 1A6A), DRB5*0101 (PDB ID: 1H15), DRB3*0101 (PDB ID: 2Q6W), DRB1*0401 (PDB ID: 2SEB), and DRB1*0301 (PDB ID: 3C5J) alleles. Moreover, it has been proved that TLR-8 is responsible for mediating the immune responses against the RNA viruses and TLR-3 is responsible for mediating immune responses against the DNA viruses [151][152]. The coronavirus is an RNA virus [153]. For this reason, the vaccine constructs of Wuhan novel coronavirus were docked against TLR-8 (PDB ID: 3W3M). The protein-protein docking study was conducted using different online docking tools to improve the prediction accuracy of the docking study. At first, the docking was performed by ClusPro 2.0 (https://cluspro.bu.edu/login.php). The server ranks the docked complexes based on their center scores and lowest energy scores [154]-[156]. The binding affinity (ΔG in kcal mol^-1^) of the docked complexes were later analyzed by PRODIGY tool of HADDOCK webserver (https://haddock.science.uu.nl/). The lower binding energy of the server represents the higher binding affinity and vice versa [157]-[159]. Next, the docking was again carried out by PatchDock (https://bioinfo3d.cs.tau.ac.il/PatchDock/php.php) server and later refined and re-scored by FireDock server (http://bioinfo3d.cs.tau.ac.il/FireDock/php.php). The FireDock server ranks the docked complexes based on their global energy and the lower global energy represents the better result. Finally, the docking was conducted using HawkDock server (http://cadd.zju.edu.cn/hawkdock/). At this stage, the Molecular Mechanics/Generalized Born Surface Area (MM-GBSA) study was also calculated by the HawkDock server which works on specific algorithm that depicts that, the lower score and lower energy represent the better scores [160]-[163]. From the docking experiment, one best vaccine was selected based on the docking score. The docked structures were visualized by PyMol tool [164].

### 2.15. Molecular Dynamic Simulation

The molecular dynamics (MD) simulation study was of the best selected vaccine construct was carried out using the online MD simulation tool, iMODS (http://imods.chaconlab.org/). It is a fast, online, user-friendly and effective molecular dynamics simulation server that predicts the deformability, B-factor (mobility profiles), eigenvalues, variance, co-variance map and elastic network of the protein complex. For a protein complex, the stability depends on the ability to deform at each of its amino acids. The eigenvalue represents the motion stiffness of the protein complex and the lower eigenvalue represents easy deformability of the complex. The server also determines and measures the protein flexibility [165]-[169].

### 2.16. Codon Adaptation and In Silico Cloning

The best predicted vaccine from the previous steps, was reverse transcribed to a possible DNA sequence which is supposed to express the vaccine protein in a target organism. The cellular machinery of that particular organism could use the codons of the newly adapted DNA sequence efficiently for producing the desired vaccine. Codon adaptation is a necessary step of vaccine design because this step provides the effective prediction of the DNA sequence of a vaccine construct. An amino acid can be encoded by different codons in different organisms, which is known as codon bias. Codon adaptation predicts the best codon for a specific amino acid that should work effectively and efficiently in a specific organism. The best predicted vaccine was used for codon adaptation by the Java Codon Adaptation Tool or JCat server (http://www.jcat.de/). The server ensures the maximal expression of protein in a target organism. Eukaryotic *E. coli* strain K12 was selected at the JCat server and rho-independent transcription terminators, prokaryotic ribosome binding sites and SgrA1 and SphI cleavage sites of restriction enzymes, were avoided. The optimized DNA sequence was then taken and SgrA1 and SphI restriction sites were attached to the N-terminal and C-terminal sites, respectively. Finally, the SnapGene restriction cloning module was used to insert the newly adapted DNA sequence between the SgrA1 and SphI restriction sites of pET-19b vector [170]-[174].

## 3. Results

### 3.1 Identification, Selection and Retrieval of Viral Protein Sequences

The SARS-CoV-2 (Wuhan seafood market pneumonia virus) was identified from the NCBI database (https://www.ncbi.nlm.nih.gov/). Four protein sequences i.e., Nucleocapsid Phosphoprotein (accession no: QHD43423.2), Membrane Glycoprotein (accession no: QHD43419.1), ORF3a Protein (accession no: QHD43417.1) and Surface Glycoprotein (accession no: QHD43416.1) were selected for possible vaccine construction and retrieved from the NCBI database in fasta format. **Table 01** lists the proteins sequences with their NCBI accession numbers.

**Table 01.** Table lists the proteins of COVID-19 used in the study with their PubMed accession numbers.

| Serial no. | Name of the protein | PubMed accession no. |
| --- | --- | --- |
| 01 | Nucleocapsid phosphoprotein | QHD43423.2 |
| 02 | Membrane glycoprotein | QHD43419.1 |
| 03 | ORF3a protein | QHD43417.1 |
| 04 | Surface glycoprotein | QHD43416.1 |

### 3.2. Antigenicity Prediction and Physicochemical Property Analysis of the Protein Sequences

Two proteins: nucleocapsid phosphoprotein and surface glycoprotein, were identified as potent antigens and hence used in the next phases of the experiment (**Table 02**). The physicochemical property analysis was conducted for these two selected proteins. Nucleocapsid phosphoprotein had the highest predicted theoretical pI of 10.07, however, surface glycoprotein had the highest predicted extinction co-efficient of 148960 M^-1^ cm^-1^. Both of them were found to have similar predicted half-life of 30 hours. However, surface glycoprotein had the highest predicted aliphatic index and grand average of hydropathicity (GRAVY) values among the two proteins (**Table 03**). However, numerous *in vitro* and *in vivo* researches should be carried out on these predictions to determine the degree of their accuracy.

**Table 02.**
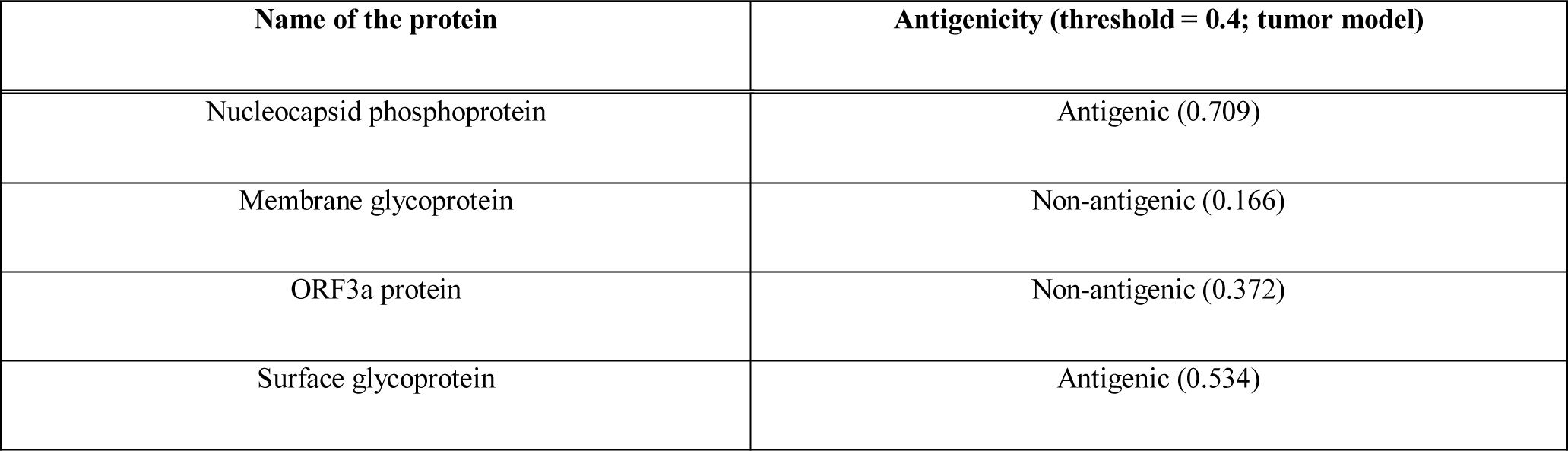
The antigenicity determination of the nine selected proteins.

| Name of the protein | Antigenicity (threshold = 0.4; tumor model) |
| --- | --- |
| Nucleocapsid phosphoprotein | Antigenic (0.709) |
| Membrane glycoprotein | Non-antigenic (0.166) |
| ORF3a protein | Non-antigenic (0.372) |
| Surface glycoprotein | Antigenic (0.534) |

**Table 03.** The antigenicity and physicochemical property analysis of the selected viral proteins.

| Name of the protein sequence | Total amino acids | Molecular weight | Theoretical pI | Ext. coefficient (in $M^{-1} cm^{-1}$) | Est. half-life (in mammalian cell) | Aliphatic index | Grand average of hydropathicity (GRAVY) |
| --- | --- | --- | --- | --- | --- | --- | --- |
| Nucleocapsid phosphoprotein | 419 | 45625.70 | 10.07 | 43890 | 30 hours | 52.53 | -0.971 |
| Surface glycoprotein | 1273 | 141178.47 | 6.24 | 148960 | 30 hours | 84.67 | -0.079 |

### 3.3. T-cell and B-cell Epitope Prediction and their Antigenicity, Allergenicity and Topology Determination

The MHC class-I and MHC class-II epitopes, determined for potential vaccine construction. The IEDB (https://www.iedb.org/) server generates a good number of epitopes. However, based on the antigenicity scores, ten epitopes were selected from the top twenty epitopes because the epitopes generated almost similar AS and percentile scores. Later, the epitopes with high antigenicity, non-allergenicity and non-toxicity were selected for vaccine construction. The B-cell epitopes were also selected based on their antigenicity, non-allergenicity and length (the sequences with more than 10 amino acids).

**Table 04** and **Table 05** list the potential T-cell epitopes of nucleocapsid phosphoprotein and **Table 06** and **Table 07** list the potential T-cell epitopes of surface glycoprotein. **Table 08** lists the predicted B-cell epitopes of the two proteins and **Table 09** lists the epitopes that followed the mentioned criteria and were selected for further analysis and vaccine construction.

**Table 04.**
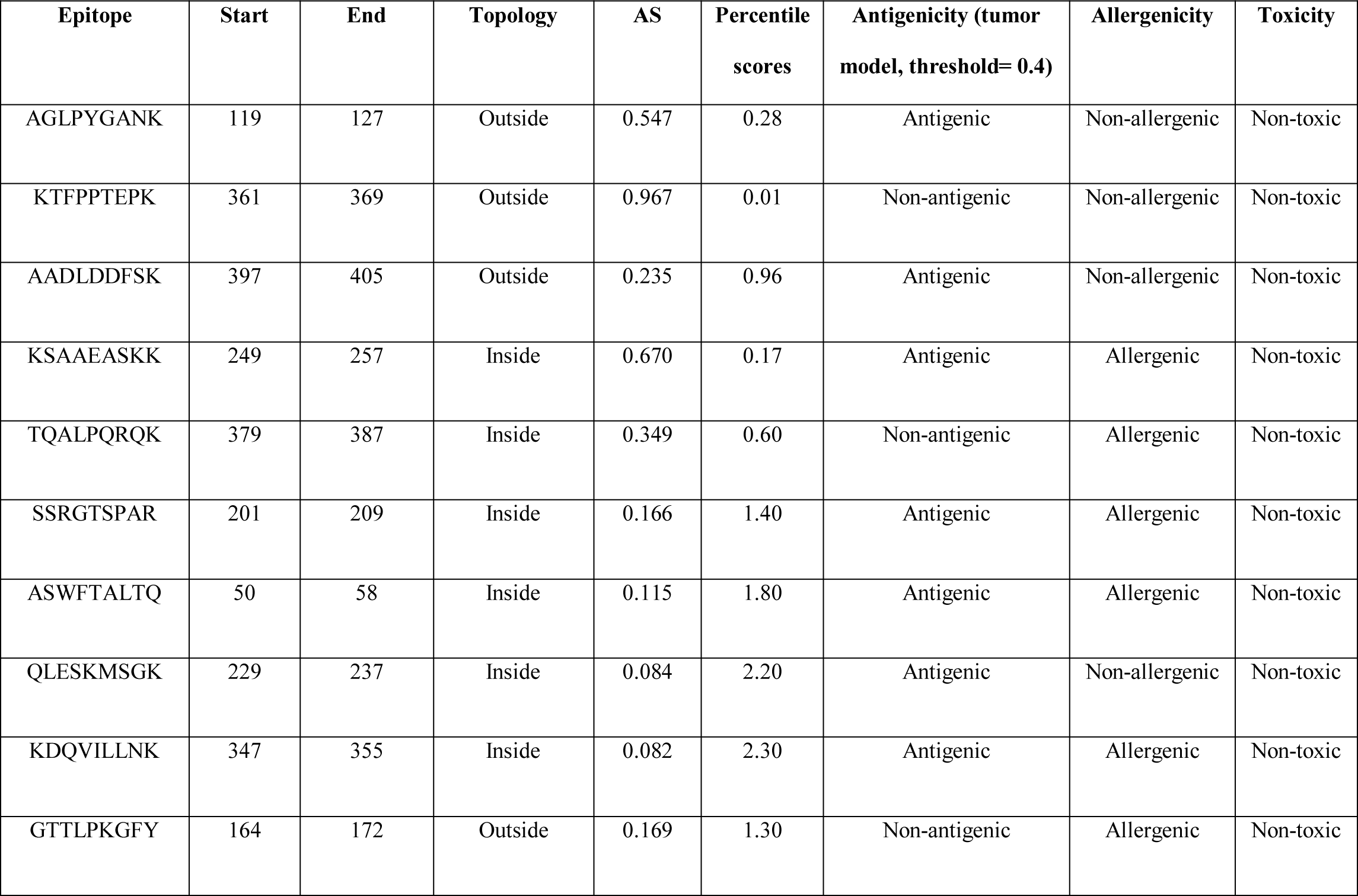
MHC class-I epitope prediction and topology, antigenicity, allergenicity and toxicity analysis of the epitopes of nucleocapsid phosphoprotein. AS: Antigenic Score.

**Table 05.** MHC class-II epitope prediction and topology, antigenicity, allergenicity and toxicity analysis of the epitopes of nucleocapsid phosphoprotein. AS: Antigenic Score.

| Epitope | Start | End | Topology | AS | Percentile scores | Antigenicity (tumor model, threshold= 0.4) | Allergenicity | Toxicity |
| --- | --- | --- | --- | --- | --- | --- | --- | --- |
| QELIRQGTDYKH | 289 | 300 | Inside | 2.90 | 3.60 | Antigenic | Non-allergenic | Non-toxic |
| ELIRQGTDYKHW | 290 | 301 | Inside | 2.90 | 3.60 | Antigenic | Allergenic | Non-toxic |
| DQELIRQGTDYK | 288 | 299 | Inside | 2.90 | 3.60 | Antigenic | Allergenic | Non-toxic |
| SRIGMEVTPSGT | 318 | 329 | Inside | 2.60 | 4.80 | Non-antigenic | Non-allergenic | Non-toxic |
| LIRQGTDYKHWP | 291 | 302 | Inside | 2.90 | 3.60 | Antigenic | Non-allergenic | Non-toxic |
| RLNQLESKMSGK | 226 | 237 | Inside | 2.00 | 8.10 | Antigenic | Non-allergenic | Non-toxic |
| WFTALTQHGKED | 52 | 63 | Inside | 1.70 | 11.0 | Antigenic | Allergenic | Non-toxic |
| LNQLESKMSGKG | 227 | 238 | Outside | 2.00 | 8.10 | Antigenic | Non-allergenic | Non-toxic |
| LDRLNQLESKMS | 224 | 235 | Inside | 2.00 | 8.10 | Antigenic | Non-allergenic | Non-toxic |
| WFTALTQHG | 49 | 60 | Outside | 1.70 | 11.0 | Antigenic | Allergenic | Non-toxic |

**Table 06.** MHC class-I epitope prediction and topology, antigenicity, allergenicity and toxicity analysis of the epitopes of surface glycoprotein. AS: Antigenic Score.

| Epitope | Start | End | Topology | AS | Percentile scores | Antigenicity (tumor model, threshold= 0.4) | Allergenicity | Toxicity |
| --- | --- | --- | --- | --- | --- | --- | --- | --- |
| GVYFASTEK | 89 | 97 | Inside | 0.938 | 0.01 | Non-antigenic | Non-allergenic | Non-toxic |
| ASANLAATK | 1020 | 1028 | Inside | 0.911 | 0.01 | Non-antigenic | Allergenic | Non-toxic |
| SVLNDILSR | 975 | 983 | Inside | 0.849 | 0.04 | Antigenic | Non-allergenic | Non-toxic |
| GVLTESNKK | 550 | 558 | Inside | 0.731 | 0.12 | Antigenic | Non-allergenic | Non-toxic |
| SSTASALGK | 939 | 947 | Outside | 0.779 | 0.09 | Antigenic | Allergenic | Non-toxic |
| GTHWFVTQR | 1099 | 1107 | Inside | 0.776 | 0.09 | Non-antigenic | Allergenic | Non-toxic |
| EILPVSMTK | 725 | 733 | Inside | 0.773 | 0.09 | Non-antigen | Allergen | Non-toxic |
| ALDPLSETK | 292 | 300 | Outside | 0.679 | 0.16 | Antigenic | Allergenic | Non-toxic |
| RLFRKSNLK | 454 | 462 | Inside | 0.677 | 0.16 | Antigenic | Non-allergenic | Non-toxic |
| QIAPGQTGK | 409 | 417 | Inside | 0.674 | 0.16 | Antigenic | Non-allergenic | Non-toxic |

**Table 07.** MHC class-II epitope prediction and topology, antigenicity, allergenicity and toxicity analysis of the epitopes of surface glycoprotein. AS: Antigenic Score.

| Epitope | Start | End | Topology | AS | Percentile scores | Antigenicity (tumor model, threshold= 0.4) | Allergenicity | Toxicity |
| --- | --- | --- | --- | --- | --- | --- | --- | --- |
| FLGVYYHKNNKS | 140 | 151 | Inside | 4.40 | 0.45 | Non-antigenic | Allergenic | Non-toxic |
| TSNFRVQPTESI | 315 | 326 | Inside | 5.30 | 0.08 | Antigenic | Non-allergenic | Non-toxic |
| VYYHKNNKSWME | 143 | 154 | Inside | 4.40 | 0.45 | Non-antigenic | Allergenic | Non-toxic |
| NFRVQPTESIVR | 317 | 328 | Inside | 5.30 | 0.08 | Antigenic | Allergenic | Non-toxic |
| GVFVSNGTHWFV | 1093 | 1104 | Outside | 4.10 | 0.78 | Non-antigenic | Allergenic | Non-toxic |
| SNFRVQPTESIV | 316 | 327 | Inside | 5.30 | 0.08 | Antigenic | Non-allergenic | Non-toxic |
| LLIVNNATNVVI | 117 | 128 | Inside | 4.30 | 0.58 | Antigenic | Non-allergenic | Non-toxic |
| EGVFVSNGTHWF | 1092 | 1103 | Outside | 4.10 | 0.78 | Non-antigenic | Allergenic | Non-toxic |
| VFVSNGTHWFVT | 1094 | 1105 | Outside | 4.10 | 0.78 | Non-antigenic | Non-allergenic | Non-toxic |
| IVNNATNVVIKV | 119 | 130 | Inside | 4.30 | 0.58 | Antigenic | Allergenic | Non-toxic |

**Table 08.**
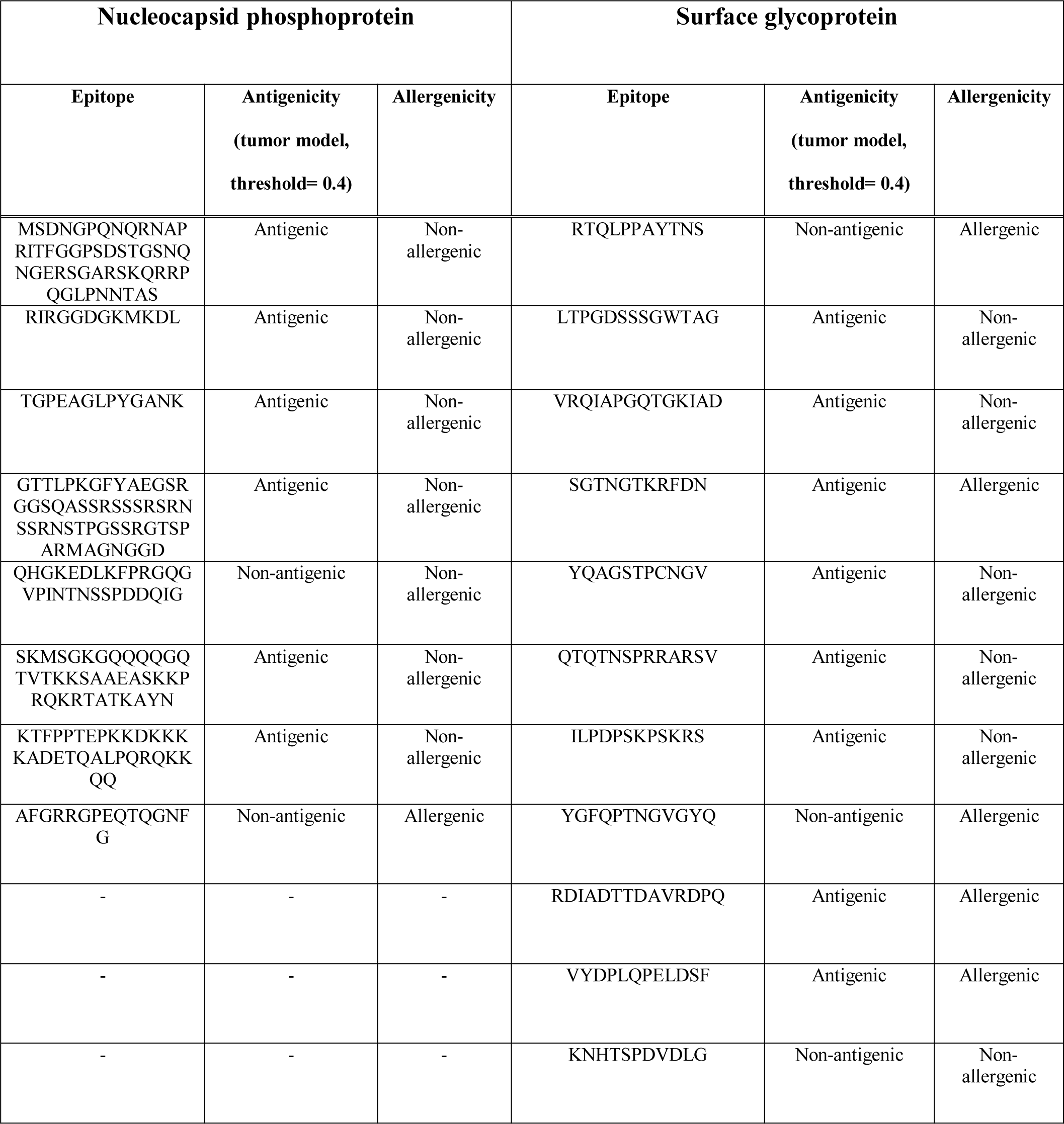
B-cell epitope prediction and topology, antigenicity, allergenicity and toxicity analysis of the epitopes of nucleocapsid phosphoprotein and surface glycoprotein.

**Table 09.**
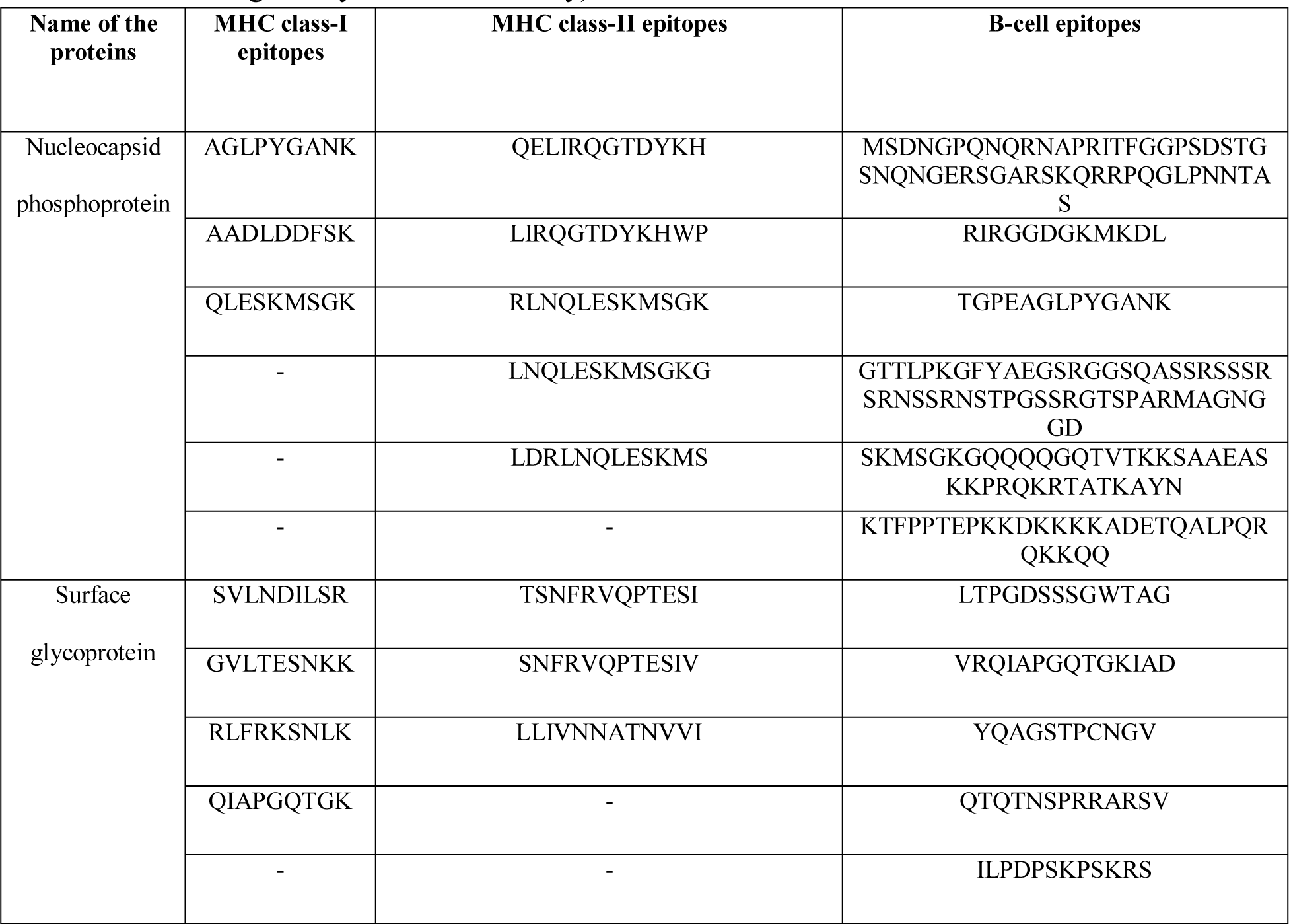
List of the epitopes that followed the selection criteria (high antigenicity, non-allergenicity and non-toxicity) and selected for vaccine construction.

### 3.4. Cluster Analysis of the MHC Alleles

The online tool MHCcluster 2.0 (http://www.cbs.dtu.dk/services/MHCcluster/), was used for the prediction or cluster analysis of the possible MHC class-I and MHC class-II alleles that may interact with the selected epitopes during the immune responses. The tool illustrates the relationship of the clusters of the alleles in phylogenetic manner. **Figure 03** depicts the result of the cluster analysis where the red zone indicates strong interaction and the yellow zone corresponds to weaker interaction.

**Figure 03.**
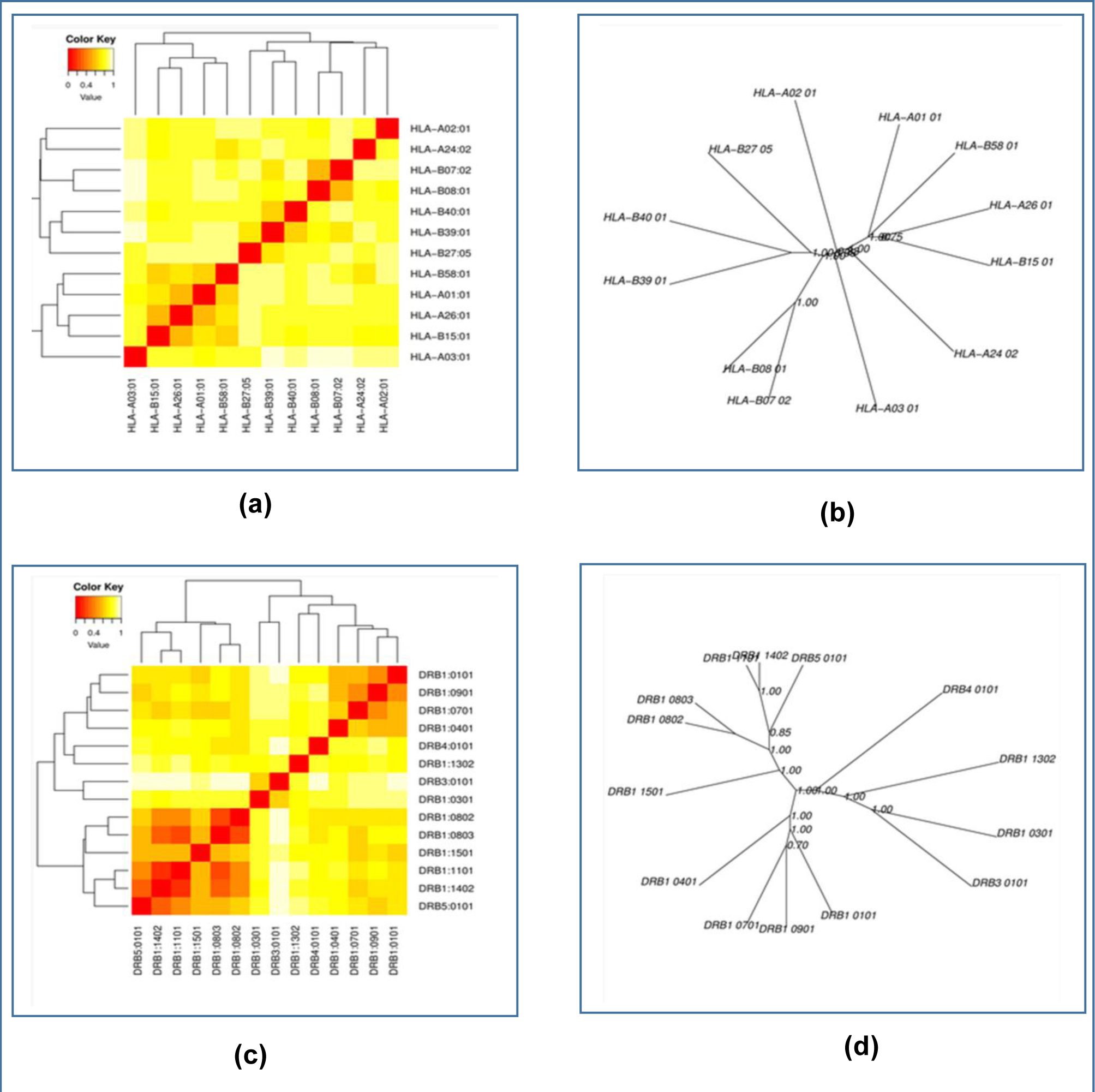
The results of the MHC cluster analysis. Here, (a) is the heat map of MHC class-I cluster analysis, (b) is the tree map of MHC class-I cluster analysis, (c) is the heat map of MHC class-II cluster analysis, (d) is the tree map of MHC class-II cluster analysis.

### 3.5. Generation of the 3D Structures of the Epitopes and Peptide-Protein Docking

After 3D structure prediction of the selected epitopes, the peptide-protein docking was conducted to find out, whether all the epitopes had the ability to bind with the MHC class-I as well as MHC class-II molecules or not. The HLA-A*11-01 allele (PDB ID: 5WJL) was used as the receptor for docking with the MHC class-I epitopes and HLA-DRB1*04-01 (PDB ID: 5JLZ) was used as the receptor for docking with the MHC class-II epitopes. Among the MHC class-I epitopes of nucleocapsid phosphoprotein, QLESKMSGK showed the best result with the lowest global energy of -53.28. Among the MHC class-II epitopes of nucleocapsid phosphoprotein, LIRQGTDYKHWP generated the lowest and best global energy score of 16.44. GVLTESNKK generated the best global energy score of -34.60 of the MHC class-I epitopes of surface glycoprotein. Among the MHC class-II epitopes of surface glycoprotein, TSNFRVQPTESI generated the best global energy score of -2.28 (**Table 10 & Figure 04**).

**Figure 04.**
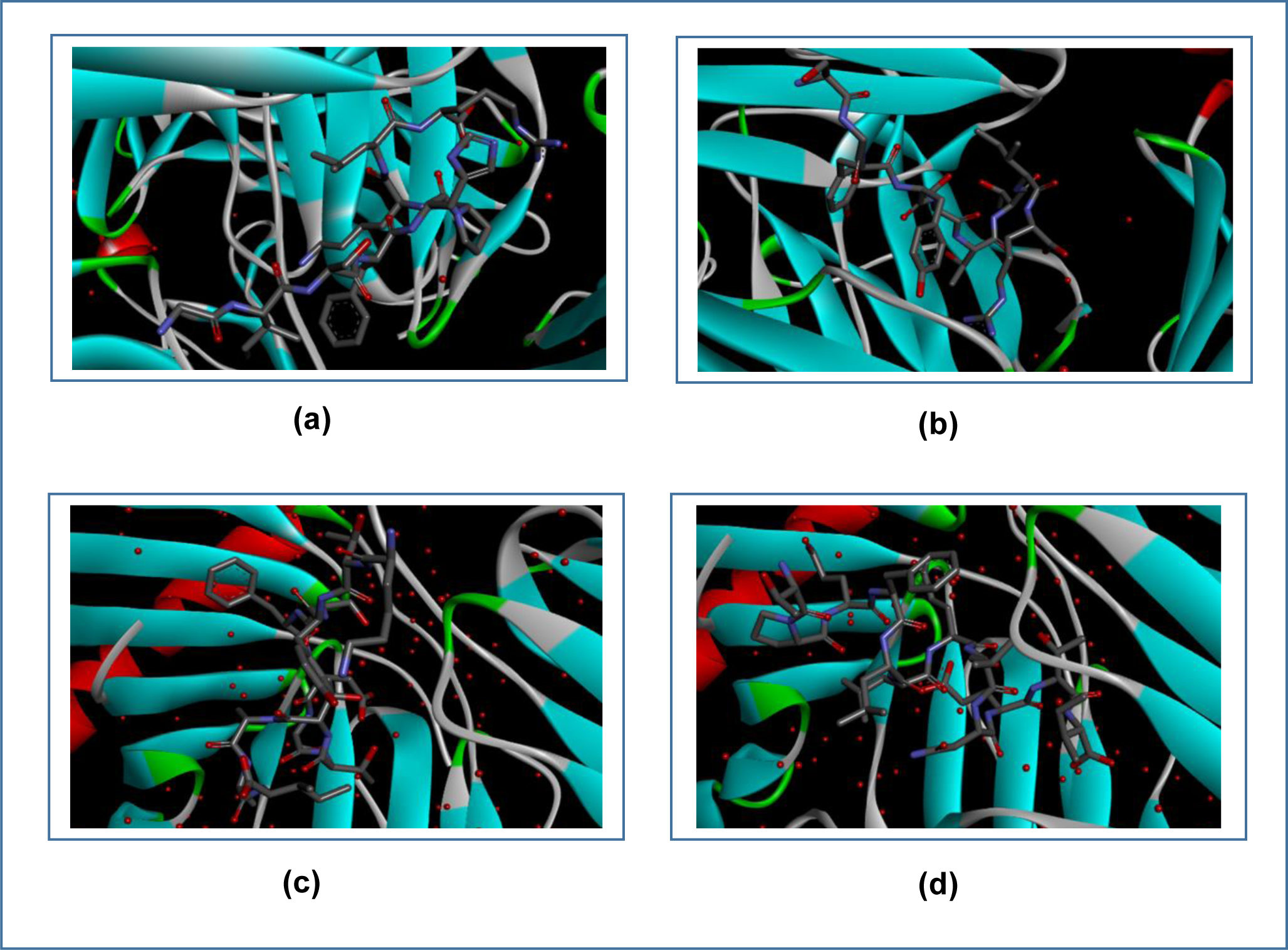
The best poses of interactions between the selected epitopes from the two proteins and their respective receptors. Here, (a) is the interaction between QLESKMSGK and MHC class-I, (b) is the interaction between GVLTESNKK and MHC class-I, (c) is the interaction between LIRQGTDYKHWP and MHC class-II, (d) is the interaction between TSNFRVQPTESI and MHC class-II. The interactions were visualized by Discovery Studio Visualizer.

**Table 10.** Results of molecular docking analysis of the selected epitopes.

| Name of the protein | Epitope | MHC allele | Global energy | Hydrogen bond energy | Epitope | MHC allele | Global energy | Hydrogen bond energy |
| --- | --- | --- | --- | --- | --- | --- | --- | --- |
| Nucleocapsid phosphoprotein | AGLPYGANK | HLA-A*11-01 allele<br>(PDB ID: 5WJL) | -38.28 | -1.53 | QELIRQG TDYKH | HLA DRB1*04-01<br>(PDB ID: 5JLZ) | 14.62 | -1.09 |
|  | AADLDDFSK |  | -15.60 | -2.69 | LIRQGTD YKHWP |  | -16.44 | -1.66 |
|  | QLESKMSGK |  | -53.28 | -4.32 | RLNQLES KMSGK |  | -12.34 | -3.41 |
|  | - |  | - | - | LNQLESK MSGKG |  | 14.37 | -9.20 |
|  | - |  | - | - | LDRLNQ LESKMS |  | -1.35 | 0.00 |
| Surface glycoprotein | SVLNDILSR |  | -27.72 | -3.74 | TSNFRVQ PTESI |  | -2.28 | 0.00 |
|  | GVLTESNKK |  | -34.60 | -3.64 | SNFRVQP TESIV |  | 1.82 | 0.00 |
|  | RLFRKSNLK |  | -27.48 | -4.77 | LLIVNNA TNVVI |  | 1.38 | 0.00 |
|  | QIAPGQTGK |  | -26.86 | -0.89 | - |  | - | - |

### 3.7. Vaccine Construction

After successful docking, three vaccines were constructed using the selected epitopes which is supposed to be directed to fight against the SARS-CoV-2. To construct the vaccines, three different adjuvants were used i.e., beta defensin, L7/L12 ribosomal protein and HABA protein and different linkers i.e., EAAAK, GGGS, GPGPG and KK linkers were used at their appropriate positions. PADRE sequence is an important sequence which was used in vaccine construction. It has the capability to increase the potency of the vaccines with minimal toxicity. Moreover, PADRE sequence also improve the CTL response, thus ensuring potent immune response [175]. The newly constructed vaccines were designated as: CV-1, CV-2 and CV-3 (**Table 11**).

**Table 11.**
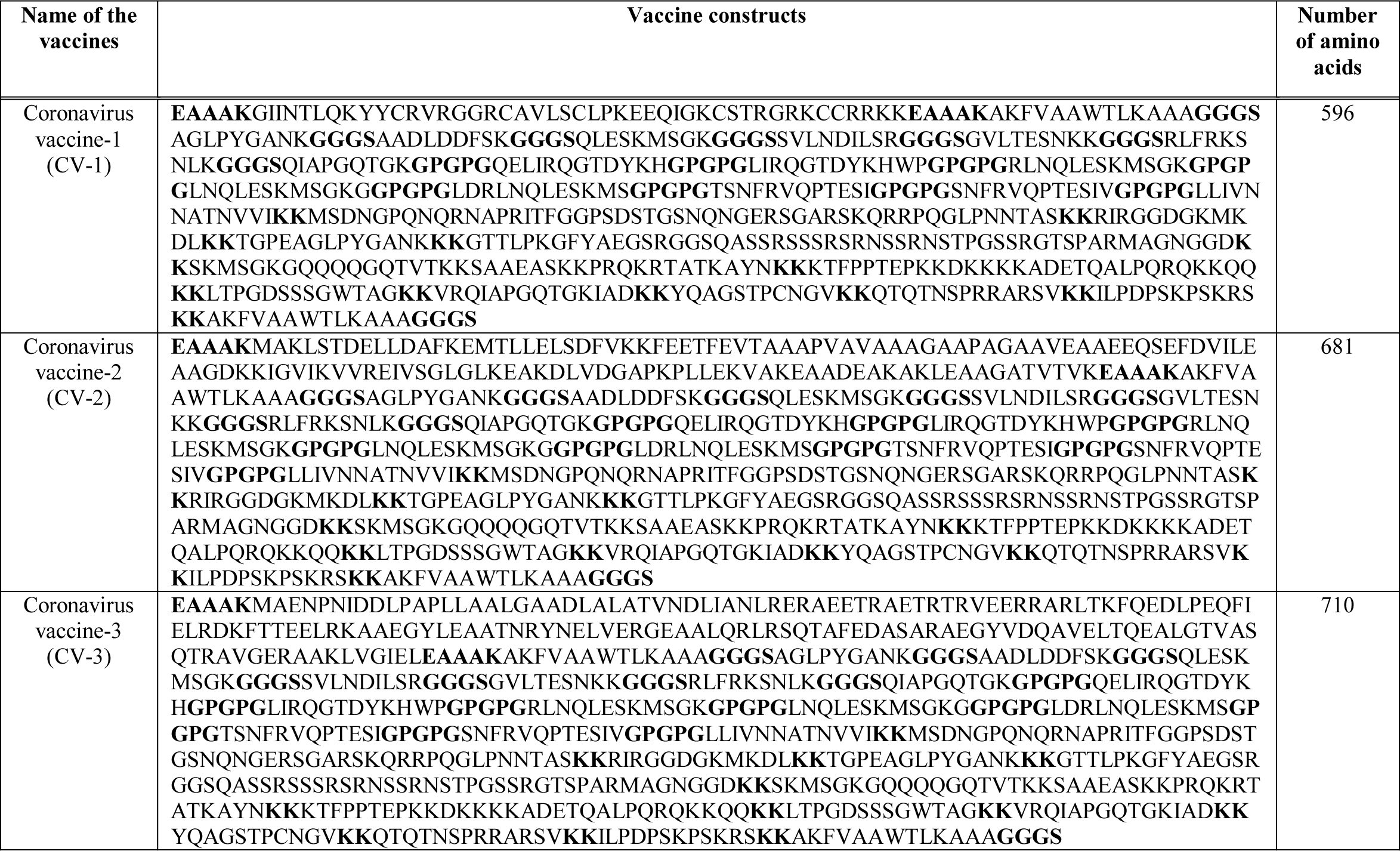
The three constructed Wuhan Novel Coronavirus vaccine constructs. In the vaccine sequences, the linkers are bolded for easy visualization.

### 3.8. Antigenicity, Allergenicity and Physicochemical Property Analysis of the Vaccine Constructs

The results of the antigenicity, allergenicity and physicochemical property analysis are listed in **Table 12**. All the three vaccine constructs were found to be antigenic as well as non-allergenic. CV-3 had the highest predicted molecular weight, extinction co-efficient and aliphatic index of 74505.61, 36900 M^-1^ cm^-1^ and 54.97 respectively. All of them had predicted *in vivo* half-life of 1 hours and CV-2 was found to possess the highest GRAVY value of -0.830 among the three vaccines.

**Table 12.** The antigenicity, allergenicity and physicochemical property analysis of the vaccine constructs. MW: Molecular Weight

| Name of the vaccine constructs | Total amino acids | Antigenicity (tumor model, threshold=0.4) | Allergenicity | MW | Theoretical pI | Ext. coefficient (in $M^{-1} cm^{-1}$) | Est. half-life (in mammalian cell) | Aliphatic index | Grand average of hydropathicity (GRAVY) |
| --- | --- | --- | --- | --- | --- | --- | --- | --- | --- |
| CV-1 | 596 | Antigenic | Non-allergenic | 62038.16 | 10.69 | 35785 | 1 hours | 46.26 | -1.041 |
| CV-2 | 681 | Antigenic | Non-allergenic | 70317.45 | 10.23 | 32430 | 1 hours | 55.86 | -0.830 |
| CV-3 | 710 | Antigenic | Non-allergenic | 74505.61 | 10.31 | 36900 | 1 hours | 54.97 | -0.941 |

### 3.9. Secondary and Tertiary Structure Prediction of the Vaccine Constructs

From the secondary structure analysis, it was determined that, the CV-1 had the highest percentage of the amino acids (67.1%) in the coil formation as well as the highest percentage of amino acids (8%) in the beta-strand formation. However, CV-3 had the highest percentage of 37.8% of amino acids in the alpha-helix formation (**Figure 05** and **Table 13**). CV-1 and CV-2 vaccines had 02 domains, whereas, CV-3 had only one domain. CV-2 had the lowest p-value of 6.35e-05. The p-value represents the relative quality of a protein model. The smaller p-value refers to higher quality of the protein model and vice-verse [176]. For this reason, CV-2 showed the best performance in the 3D structure generation experiment. Moreover, three different templates were used for generating3D structures of the three different vaccines. The RaptorX server used the templates for generating the 3D structures of the query vaccine constructs [177]. The results of the 3D structure analysis are listed in **Table 14** and illustrated in **Figure 06**.

**Figure 05.**
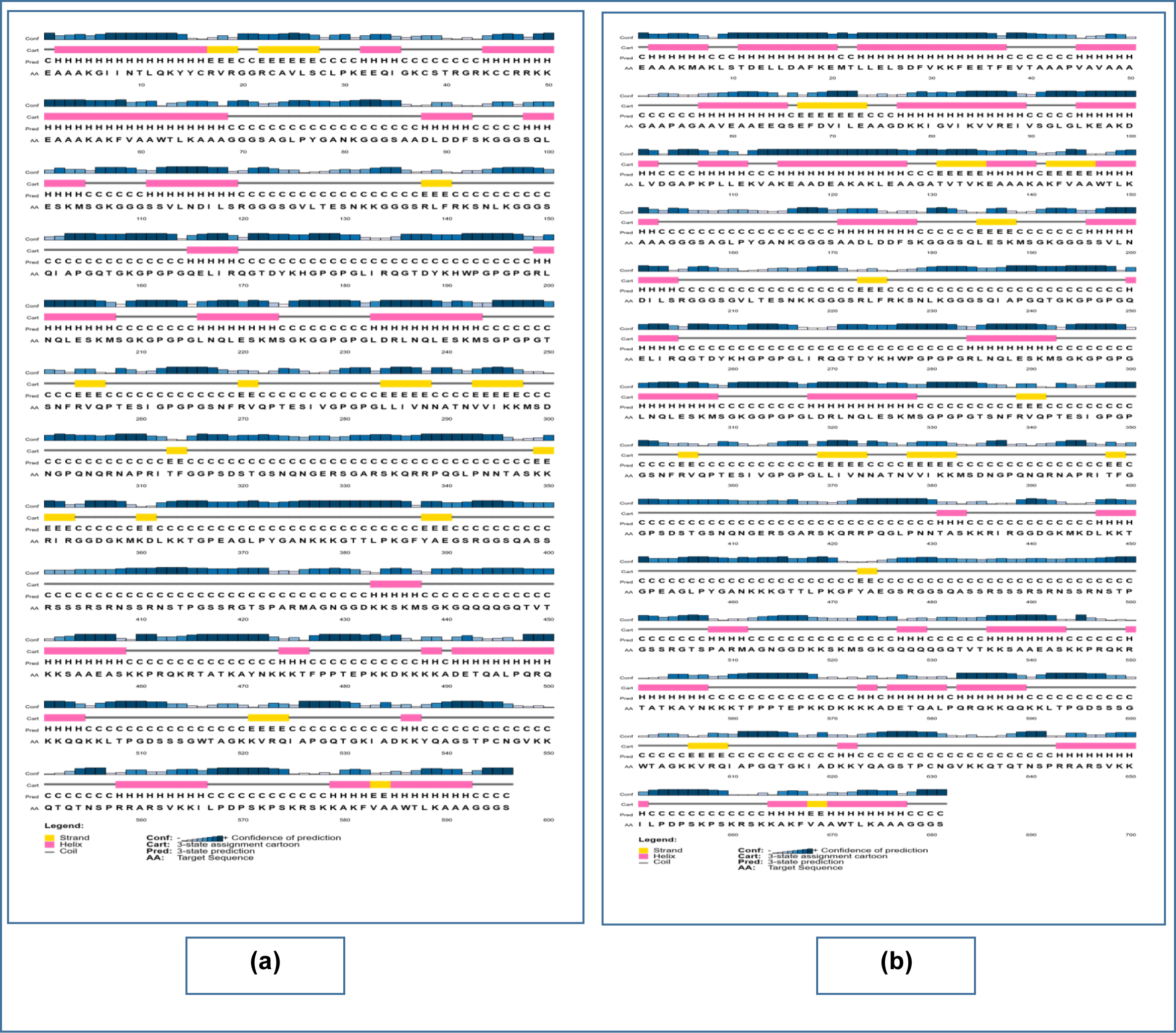
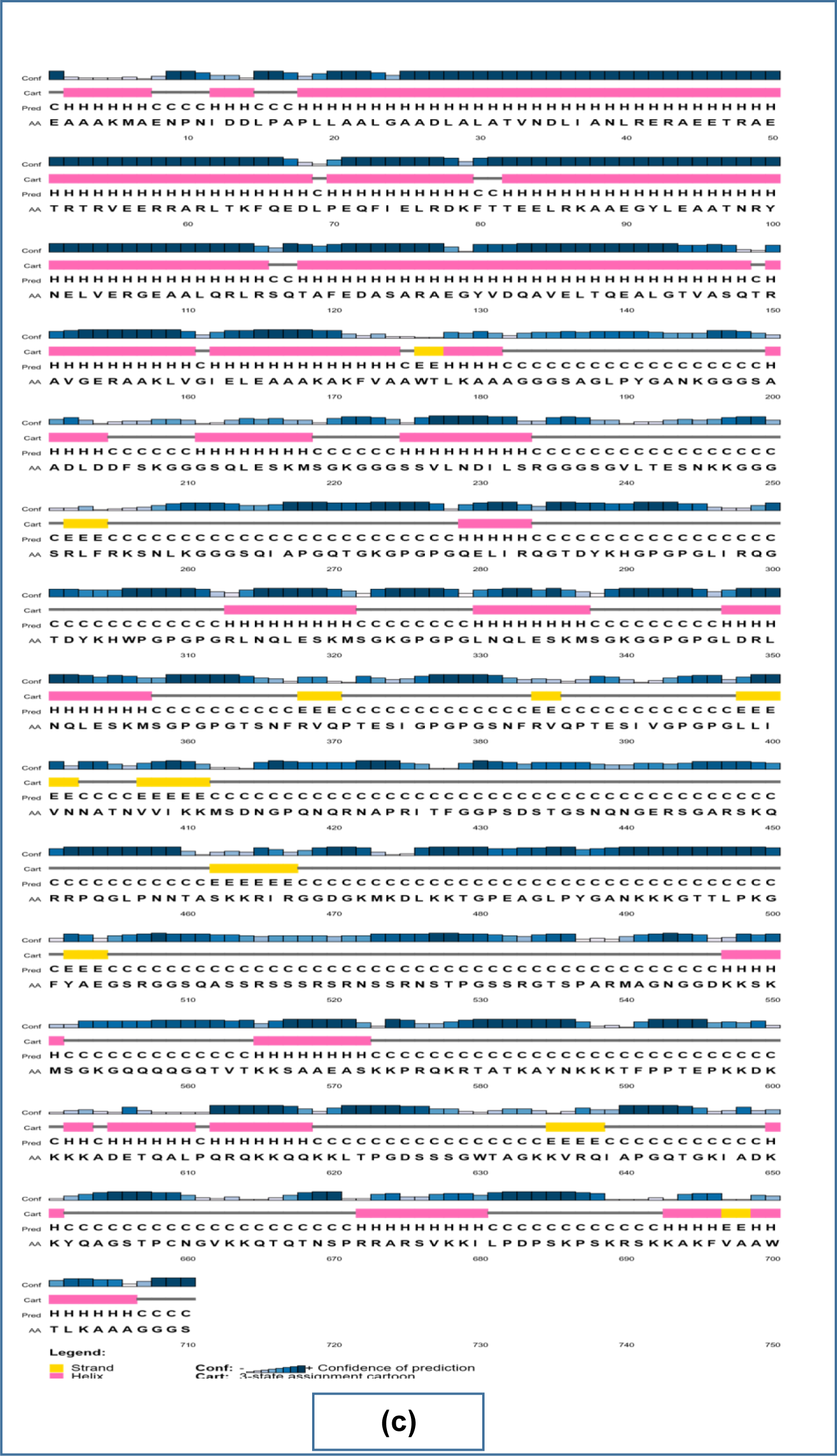
Results of the secondary structure prediction of the three vaccine constructs. Here, (a) is the CV-1 vaccine, (b) is the CV-2 vaccine, (c) is the CV-3 vaccine.

**Figure 06.**
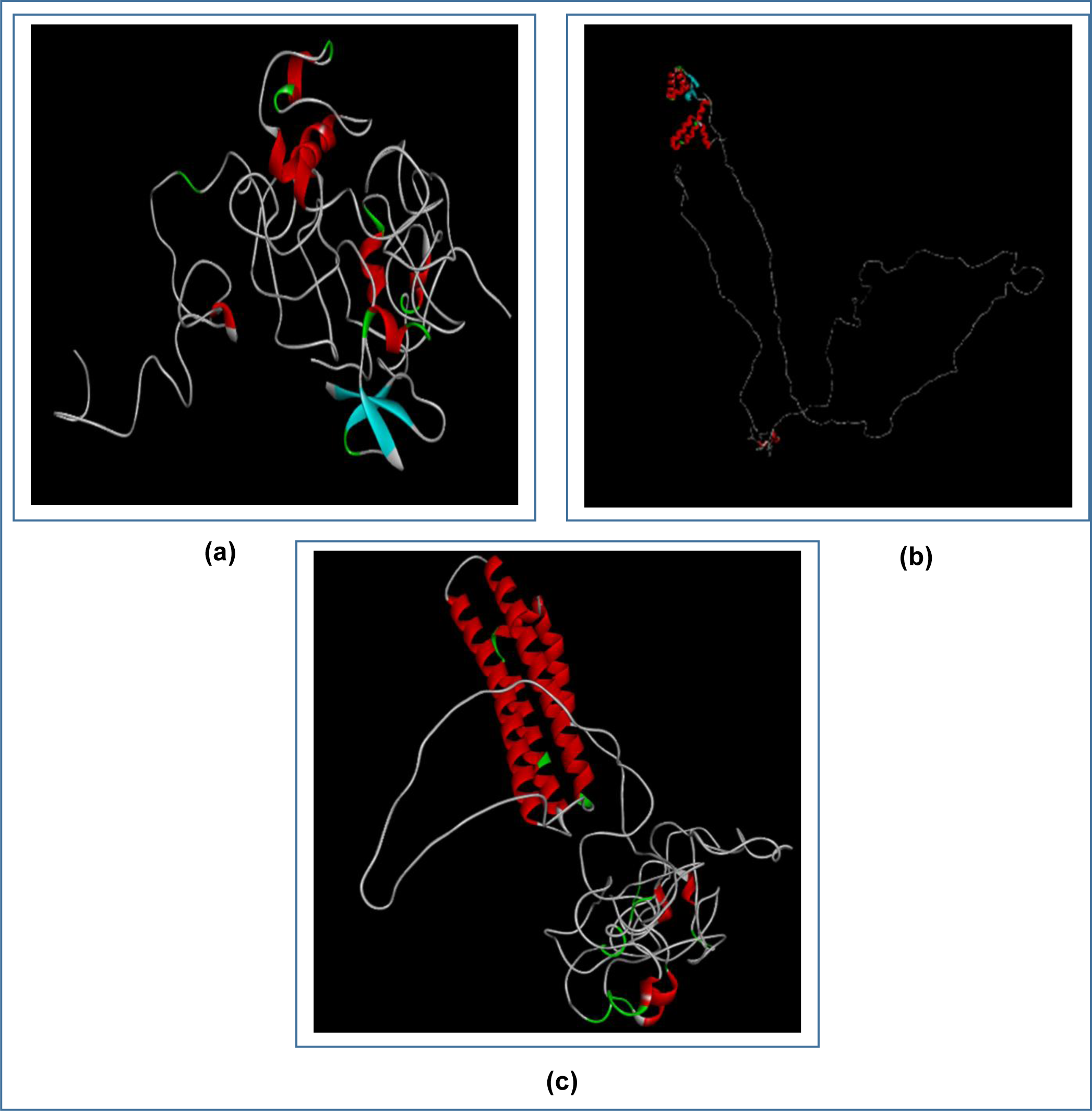
3D structures of the three predicted vaccine constructs. Here, (a) is CV-1, (b) is CV-2, (c) is CV-3.

**Table 13.** Results of the secondary structure analysis of the vaccine constructs.

| Name of the vaccine | Alpha helix (percentage of amino acids) | Beta sheet (percentage of amino acids) | Coil structure (percentage of amino acids) |
| --- | --- | --- | --- |
| CV-1 | 25% | 8% | 67.1% |
| CV-2 | 31.6% | 6.8% | 61.6% |
| CV-3 | 37.8% | 5% | 57.2% |

**Table 14.** Results of the tertiary structure analysis of the vaccine constructs.

| Name of the vaccine | Number of the domains | p-value | PDB Id of the est. matched template |
| --- | --- | --- | --- |
| CV-1 | 02 | 2.37e-04 | 1kj6A |
| CV-2 | 02 | 6.35e-05 | 1dd3A |
| CV-3 | 01 | 2.36e-04 | 6cfeA |

### 3.10. 3D Structure Refinement and Validation

The three vaccine constructs were refined and then validated in the 3D structure refinement and validation step. The PROCHECK server (https://servicesn.mbi.ucla.edu/PROCHECK/) divides the Ramachandran plot into four regions: the most favored region (represented by red color), the additional allowed region (represented by yellow color), the generously allowed region (represented by light yellow color) and the disallowed region (represented by white color). According to the server, a valid protein (the best quality protein) should have over 90% of its amino acids in the most favored region. The additional allowed region and generously allowed region might also contain some percentage of the amino acids of the protein. Moreover, no amino acid should reside in the disallowed region [178]-[180].

The 3D protein structures generated in the previous step were refined for further analysis and validation. The refined structures were refined with the aid of the Ramachandran Plots. The analysis showed that CV-1 vaccine had excellent percentage of 94.3% of the amino acids in the most favored region, 4.4% of the amino acids in the additional allowed regions, 0.0% of the amino acids in the generously allowed regions and 1.3% of the amino acids in the disallowed regions. The CV-2 vaccine had 90.0% of the amino acids in the most favored regions, 8.3% of the amino acids in the additional allowed regions, 0.6% of the amino acids in the generously allowed regions and 1.1% of the amino acids in the disallowed regions. The CV-3 vaccine showed the worst result with 77.4% of the amino acids in the most favored regions, 20.9% of the amino acids in the additional allowed regions, 1.4% of the amino acids in the generously allowed regions and 0.3% of the amino acids in the disallowed regions (**Figure 07).**

**Figure 07.**
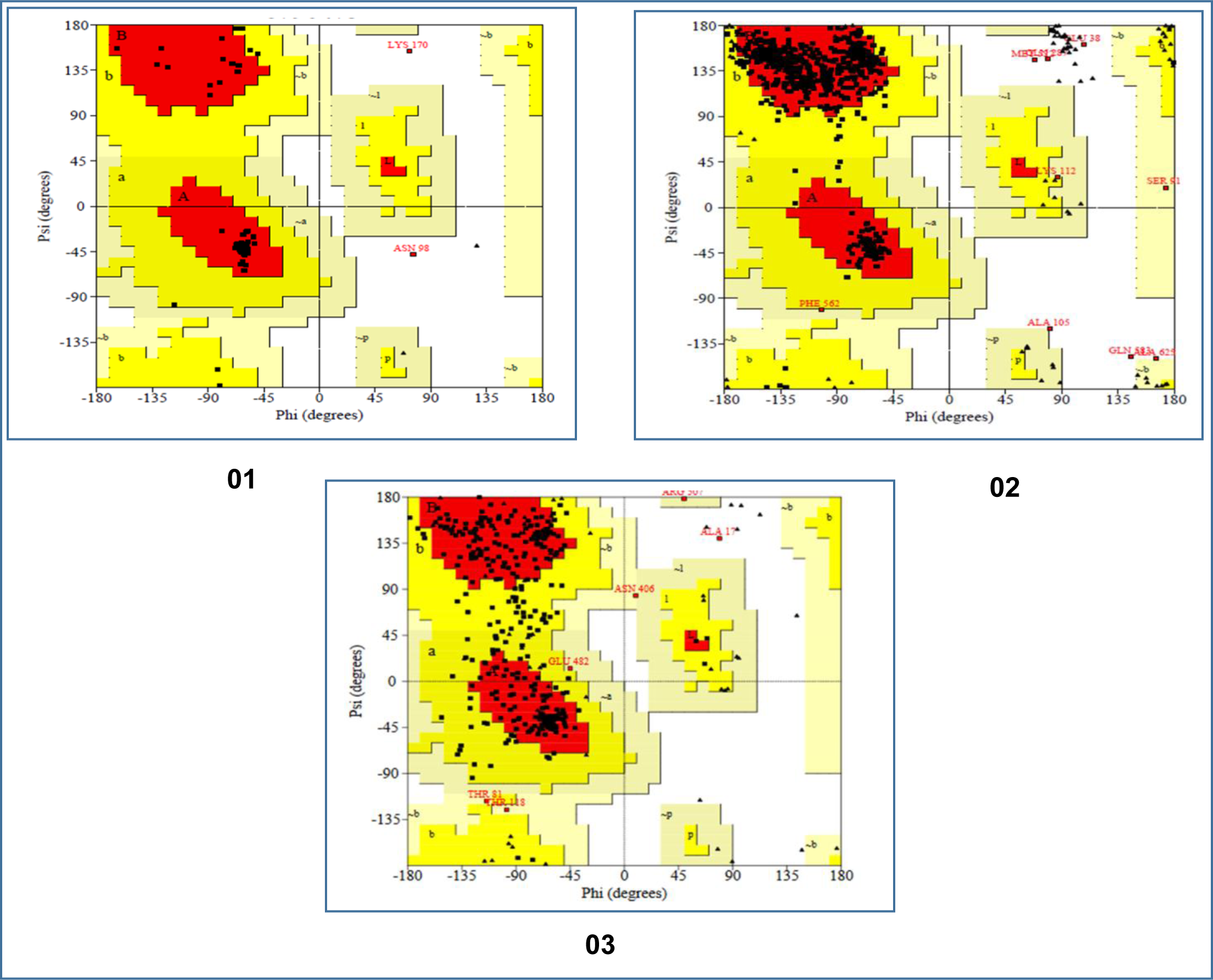
The results of the Ramachandran plot analysis of the three coronavirus vaccine constructs. Here, 01. CV-1 vaccine, 02. CV-2 vaccine, 03. CV-3 vaccine.

### 3.11. Vaccine Protein Disulfide Engineering

In protein disulfide engineering, disulfide bonds were generated within the 3D structures of the vaccine constructs. In the experiment, the amino acid pairs that had bond energy value less than 2.00 kcal/mol were selected. Since about 90% of the native disulfide bonds in proteins have energy value of less than 2.2 kcal/mol, the bond energy value of 2.00 kcal/mol was selected as the cut-off value for the experiment for better prediction [181]. The CV-1 generated 10 amino acid pairs that had the capability to form disulfide bonds. However, only one pair was selected because they had the bond energy, less than 2.00 kcal/mol: 276 Ser-311 Arg. However, CV-2 and CV-3 generated 04 and 05 pairs of amino acids, respectively, that might form disulfide bonds and no pair of amino acids showed bond energy less than 2.00 Kcal/mol. The selected amino acid pairs of CV-1 formed the mutant version of the original vaccines (**Figure 08**).

**Figure 08.**
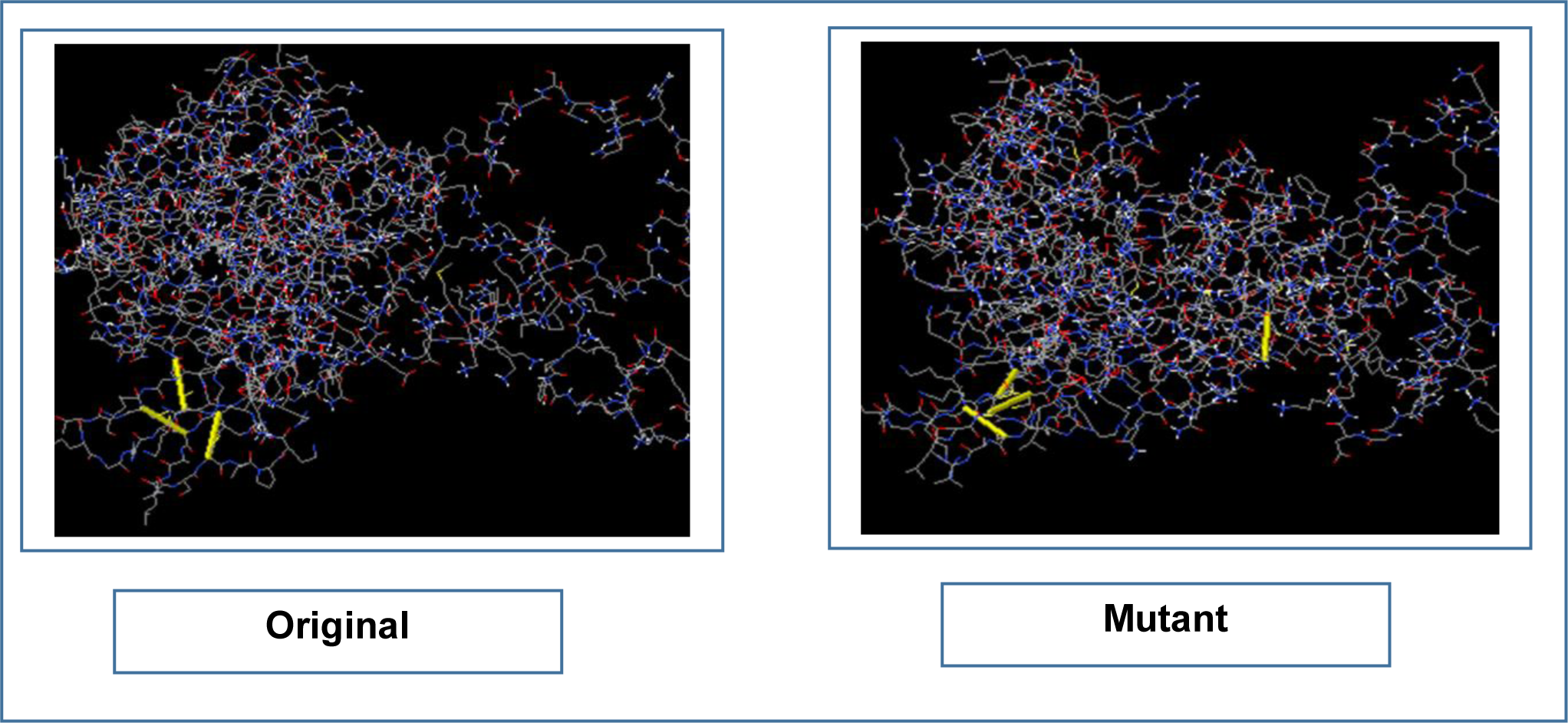
The disulfide engineering of CV-1. The original form is illustrated in the left side and the mutant form is illustrated in the right side.

### 3.12. Protein-Protein Docking Study

The protein-protein docking study was carried out to find out the best constructed COVID-19 vaccine. The vaccine construct with the best result in the molecular docking, was considered as the best vaccine construct. According to docking results, it was found that CV-1 was the best constructed vaccine. CV-1 showed the best and lowest scores in the docking as well as in the MM-GBSA study. However, CV-2 showed the best binding affinity (ΔG scores) with DRB3*0202 (-18.9 kcal/mol) and DRB1*0301 (-18.5 kcal/mol) when analyzed with ClusPro 2.0 and the PRODIGY tool of HADDOCK server. Moreover, when analyzed with PatchDock and FireDock servers, CV-3 showed best global energy scores with most of the MHC alleles i.e., DRB5*0101 (-10.70), DRB5*0101 (-19.59), DRB1*0101 (-17.46) and DRB3*0101 (-12.32). Since CV-1 showed the best results in the protein-protein docking study, it was considered as the best vaccine construct among the three constructed vaccines (**Figure 09** & **Table 15**). Later, the molecular dynamics simulation study and in silico codon adaptation studies were conducted only on the CV-1 vaccine.

**Figure 09.**
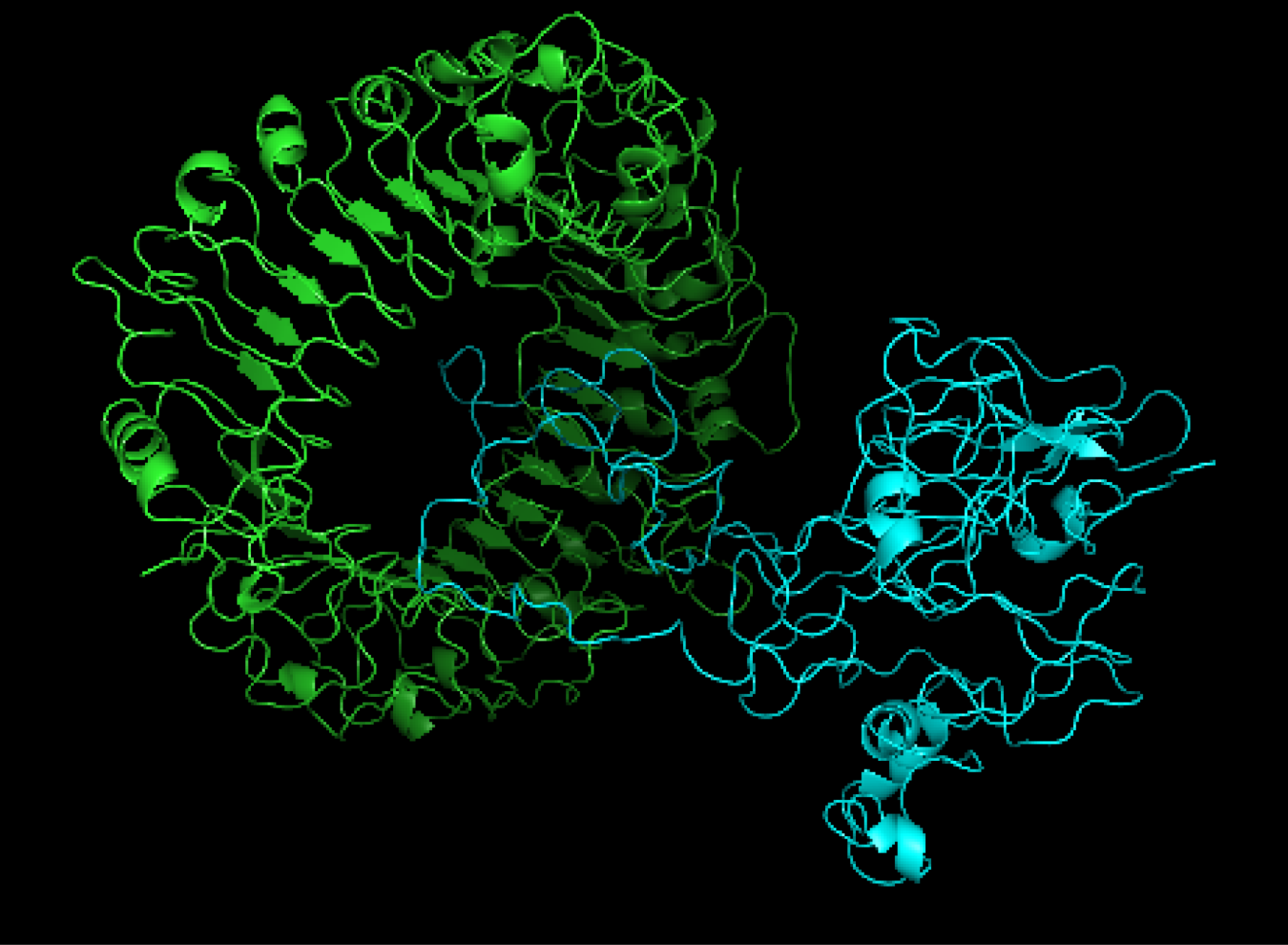
The interaction between TLR-8 (in green color) and CV-1 vaccine construct (in light blue color). The interaction was visualized with PyMol.

**Table 15.**
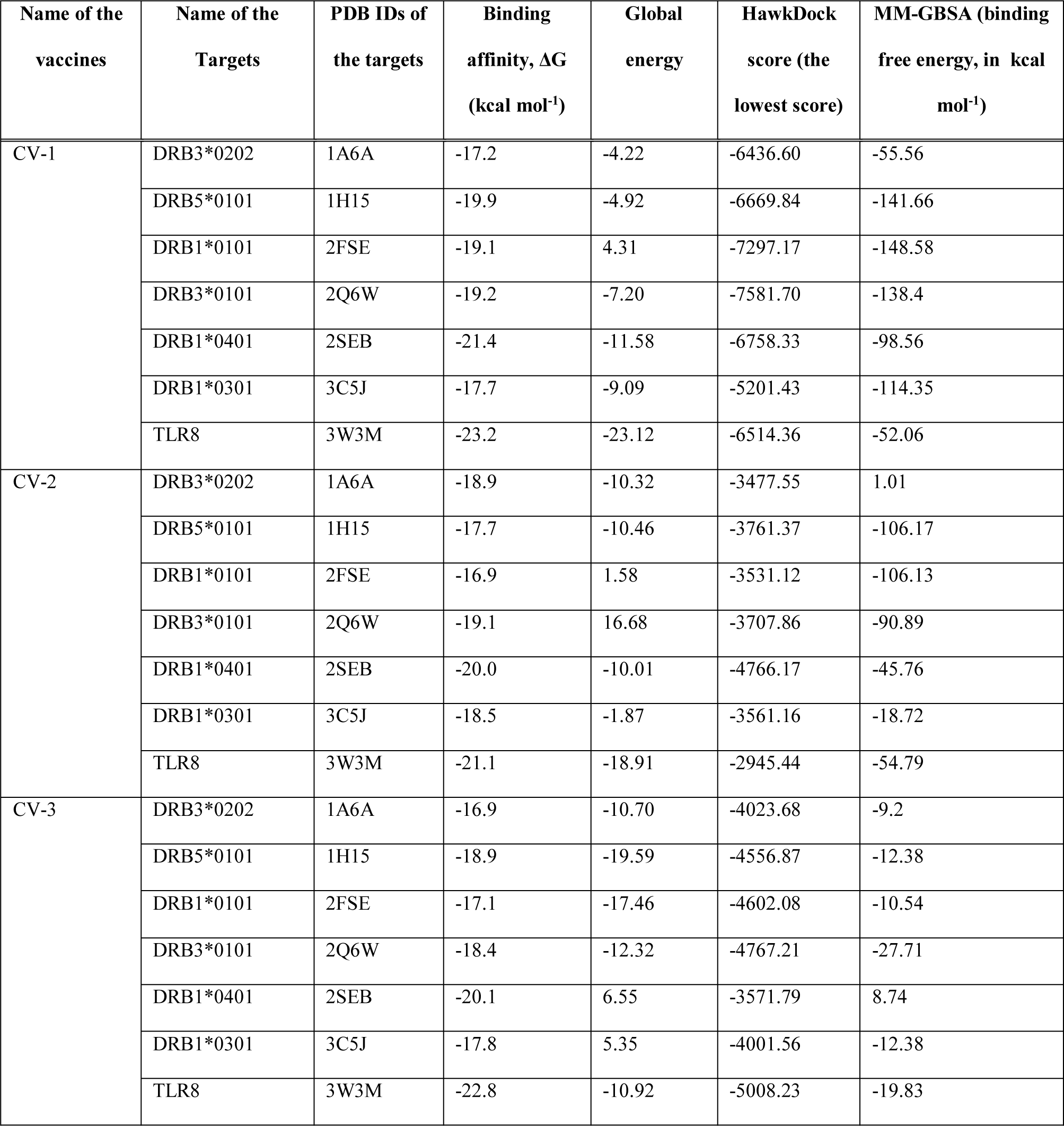
Results of the docking study of all the vaccine constructs.

### 3.13. Molecular Dynamics Simulation

The results of molecular dynamics simulation of CV-1-TLR-8 docked complex is illustrated in **Figure 10**. Dynamic simulation of proteins gives easy determination of the stability and physical movements of their atoms and molecules [182]. So, the simulation is carried out to determine the relative stability of the vaccine protein. The deformability graph of the complex illustrates the peaks representing the regions of the protein with moderate degree of deformability (**Figure 10b**). The B-factor graph of the complex gives easy visualization and comparison between the NMA and the PDB field of the docked complex (**Figure 10c)**. The eigenvalue of the docked complex is depicted in **Figure 10d**. CV-1 and TLR8 docked complex generated quite good eigenvalue of 3.817339e-06. The variance graph illustrates the individual variance by red colored bars and cumulative variance by green colored bars (**Figure 10e)**. **Figure 10f** depicts the co-variance map of the complex, where red color represents the correlated motion between a pair of residues, uncorrelated motion is indicated by white color as well as the anti-correlated motion is marked by blue color. The elastic map of the complex refers to the connection between the atoms and darker gray regions indicate stiffer regions (**Figure 09g**) [167]-[169].

**Figure 10.**
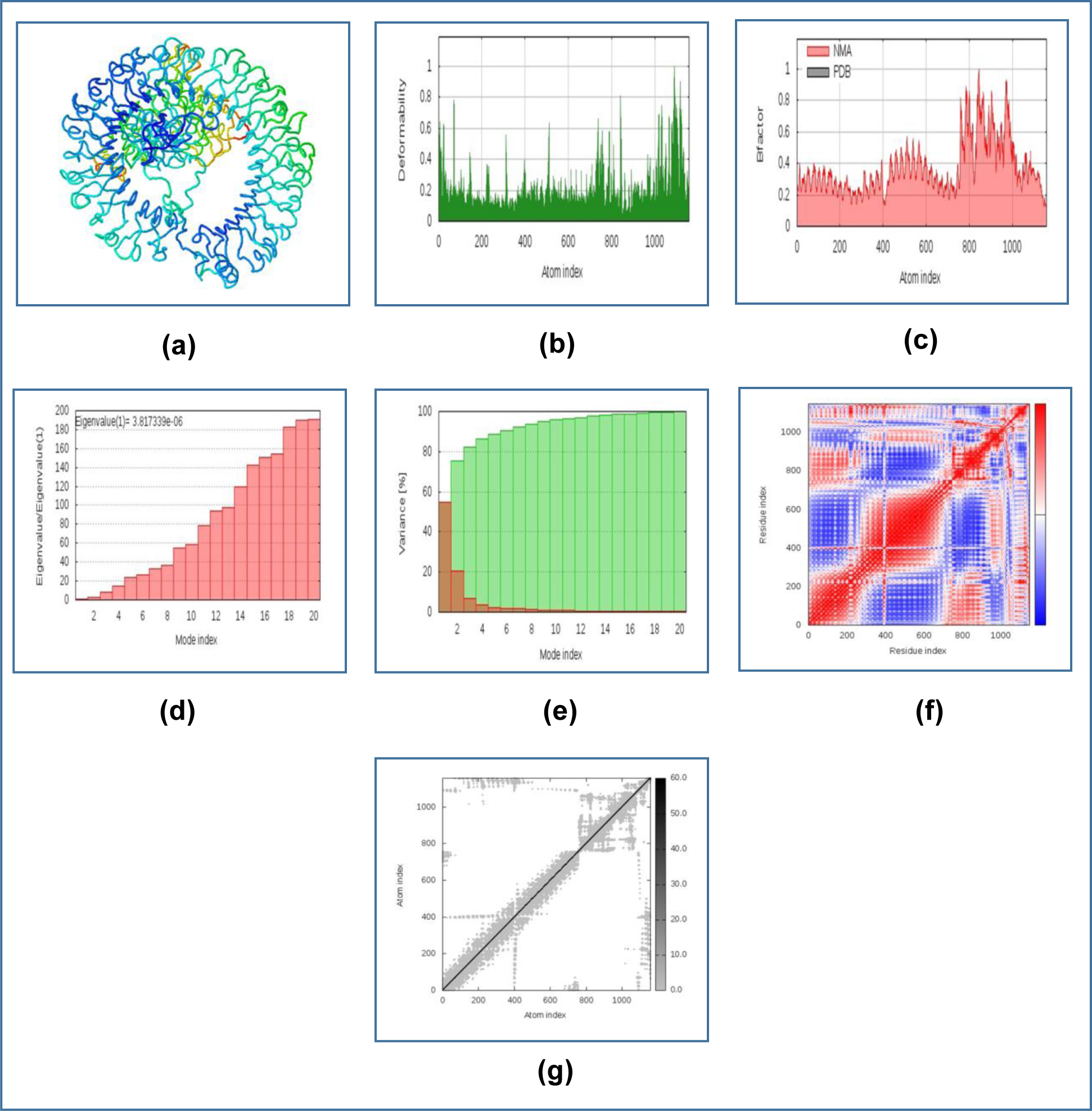
The results of molecular dynamics simulation study of CV-1 and TLR-8 docked complex. Here, (a) NMA mobility, (b) deformability, (c) B-factor, (d) eigenvalues, (e) variance (red color indicates individual variances and green color indicates cumulative variances), (f) co-variance map (correlated (red), uncorrelated (white) or anti-correlated (blue) motions) and (g) elastic network (darker gray regions indicate more stiffer regions).

### 3.14. Codon Adaptation and In Silico Cloning

Since the CV-1 protein had 596 amino acids, after reverse translation, the number nucleotides of the probable DNA sequence of CV-1 would be 1788. The codon adaptation index (CAI) value of 1.0 of CV-1 indicated that the DNA sequences contained higher proportion of the codons that should be used by the cellular machinery of the target organism *E. coli* strain K12 (codon bias). For this reason, the production of the CV-1 vaccine should be carried out efficiently [183][184]. The GC content of the improved sequence was 51.34% (**Figure 11**). The predicted DNA sequence of CV-1 was inserted into the pET-19b vector plasmid between the SgrAI and SphI restriction sites and since the DNA sequence did not have restriction sites for SgrAI and SphI restriction enzymes, SgrA1 and SphI restriction sites were conjugated at the N-terminal and C-terminal sites, respectively. The newly constructed vector is illustrated in **Figure 12**.

**Figure 11.**
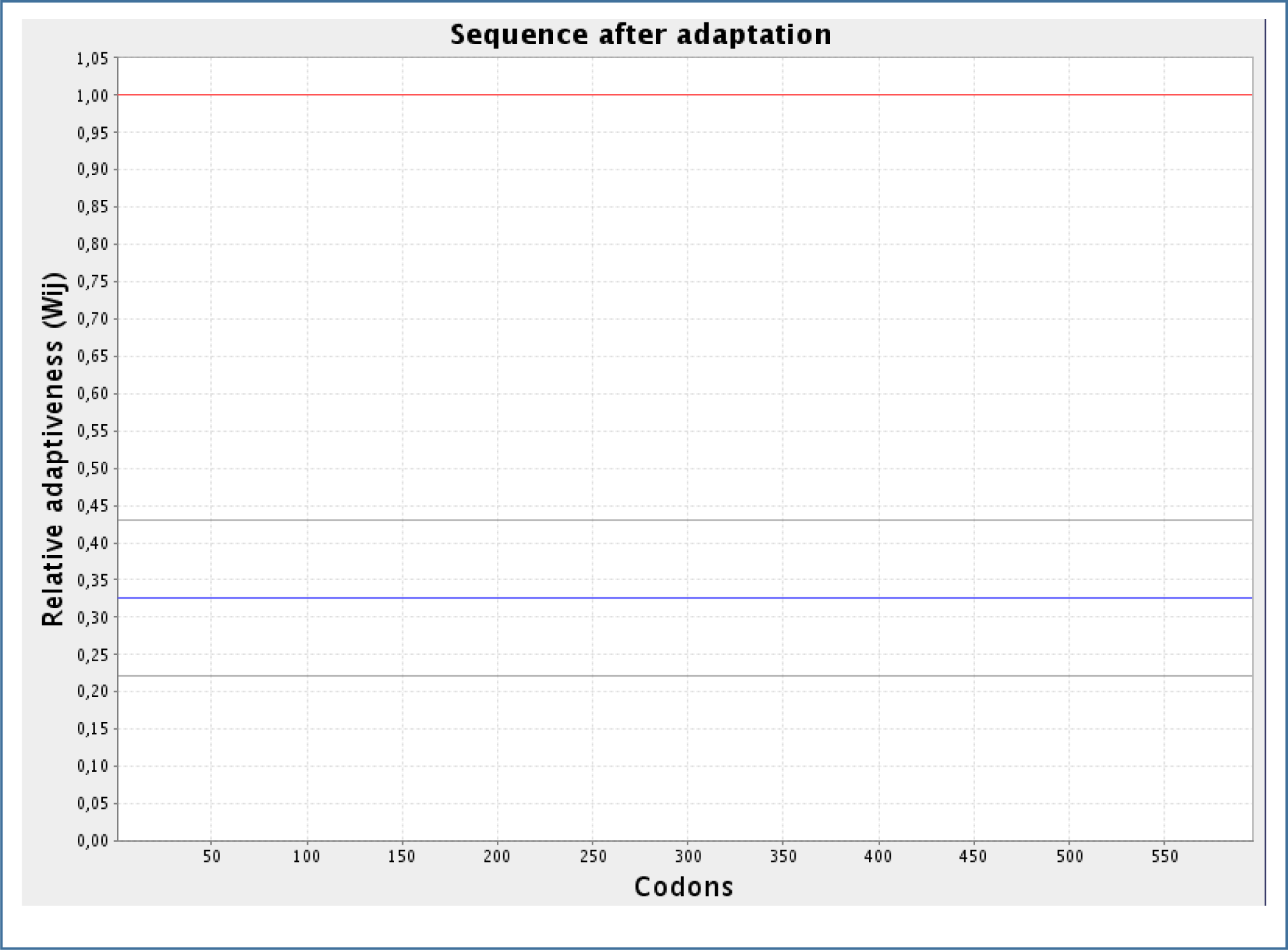
The results of the codon adaptation study of the constructed vaccine, CV-1.

**Figure 12.**
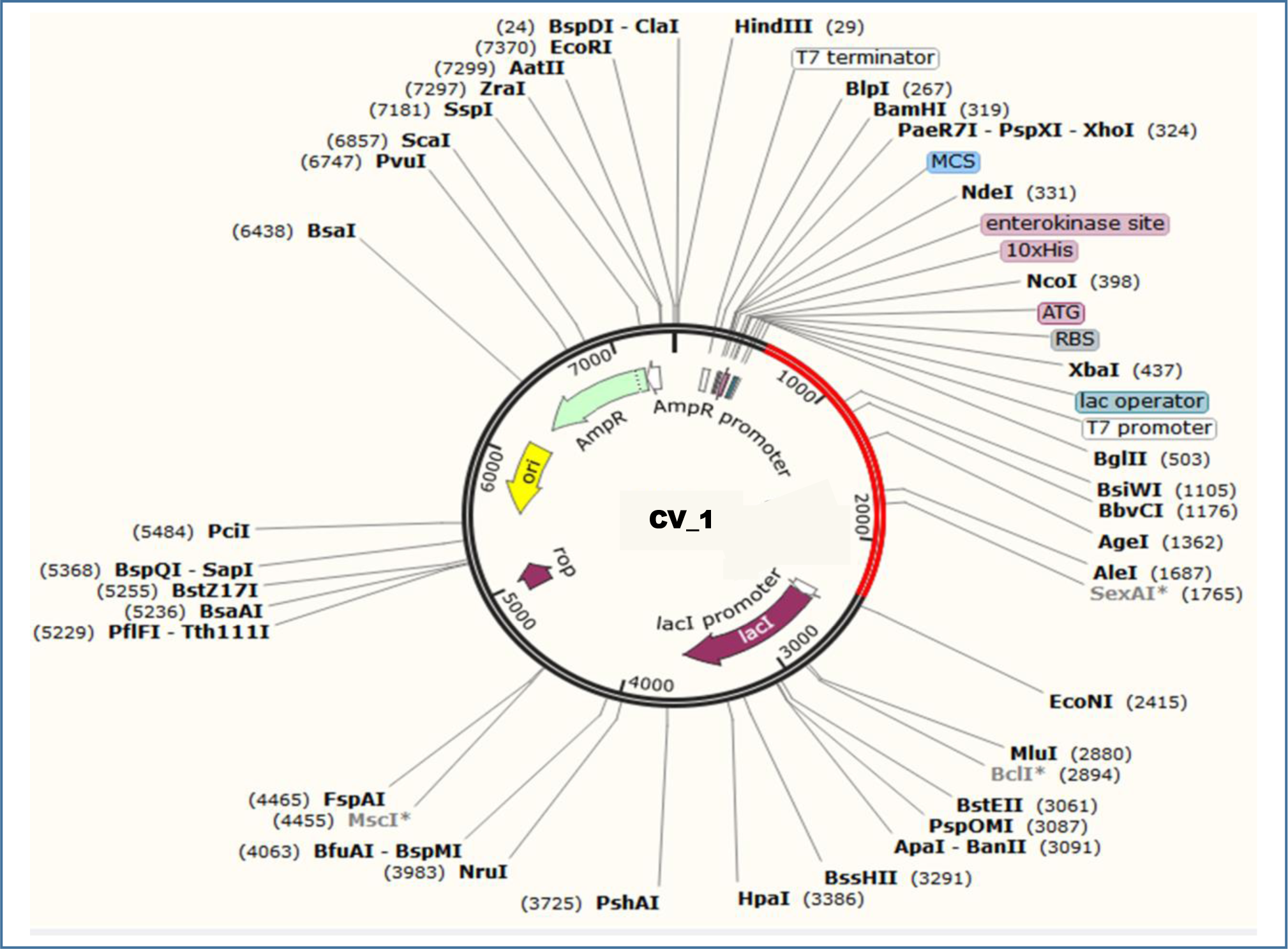
Constructed pET-19b vector with the CV-1 insert (marked in red color). In the plasmid, the larger purple colored arrow represents the *lacI* gene (from 2500 bp to 3582 bp), the smaller purple colored arrow represents the *rop* gene (from 4896 bp to 5085 bp), yellow colored arrow represents the origin of replication (from 5517 bp to 6103 bp), the light green colored arrow represents the *AmpR* (ampicillin resistance) gene (from 6274 bp to 7134 bp), the white rectangle represents the T7 terminator (from 195 bp to 242 bp), the light blue colored arrow represents the multiple cloning site (from 301 bd to 317 bp) and the desired gene has been inserted (marked by red color) between the 485 bp and 2128 bp nucleotide. Various restriction enzyme sites are mentioned in the plasmid structure.

## 4. Discussion

The current study was designed to construct possible vaccines against the Wuhan Novel Coronavirus 2019 (SARS-CoV-2), which is the cause of the recent outbreak of the deadly viral disease, COVID-19 in China. The pneumonia has already caused the death of several thousands of people worldwide. For this reason, possible vaccines were predicted in this study to fight against this lethal virus. To carry out the vaccine construction, four candidate proteins of the virus were identified and selected from the NCBI database. Only highly antigenic sequences were selected for further analysis since the highly antigenic proteins can induce better immunogenic response [185, 201]. Because the nucleocapsid phosphoprotein and surface glycoprotein were found to be antigenic, they were taken into consideration for vaccine construction.

The physicochemical property analysis was conducted for the two predicted antigenic proteins. The extinction coefficient can be defined as the amount of light that is absorbed by a particular compound at a certain wavelength [186][187]. Surface glycoprotein had the highest predicted extinction co-efficient of 148960 M^-1^ cm^-1^. The aliphatic index of a protein corresponds to the relative volume occupied by the aliphatic amino acids in the side chains of the protein, for example: alanine, valine etc. [188][189]. Surface glycoprotein also had the highest predicted aliphatic index among the two proteins (84.67). For this reason, surface glycoprotein had greater amount of aliphatic amino acids in its side chain than the nucleocapsid phosphoprotein. The grand average of hydropathicity value (GRAVY) for a protein is calculated as the sum of hydropathy values of all the amino acids of the protein, divided by the number of residues in its sequence [190]. Surface glycoprotein had the highest predicted GRAVY value of -0.079 among the two proteins. However, both of them had the predicted *in vivo* half-life of 30 hours and nucleocapsid phosphoprotein had the highest theoretical pI of 10.07. Both the proteins showed quite good results in the physicochemical property analysis.

After the physicochemical analysis of the protein sequences, the T-cell and B-cell epitope prediction was conducted. T-cell and B-cell are the two main types of cells that function in immunity. When an antigen is encountered in the body by the immune system, the antigen presenting cells or APC like macrophage, dendritic cell etc. present the antigen to the T-helper cell, through the MHC class-II molecules on their surface. The helper T-cell contains CD4+ molecule on its surface, for this reason, it is also known as CD4+ T-cell. On the other hand, other type of T-cell, cytotoxic T-cell contains CD8+ molecule on their surface, for which, they are called CD8+ T-cell. MHC class-I molecules present antigens to cytotoxic T-lymphocytes. After activation by the antigen, the T-helper cell activates the B-cell, which starts to produce large amount of antibodies. Macrophage and CD8+ cytotoxic T cell are also activated by the T-helper cell that cause the final destruction of the target antigen [191]-[195]. The possible T-cell and B-cell epitopes of the selected proteins were determined by the IEDB (https://www.iedb.org/) server. The epitopes with high antigenicity, non-allergenicity and non-toxicity were selected to vaccine construction. The B-cell epitopes (predicted by the server) that were more than ten amino acids long were taken into consideration and the antigenic and non-allergenic epitopes were selected for vaccine construction. However, most of the epitopes were found to be residing within the cell membrane.

The cluster analysis of the MHC alleles which may interact with the selected epitopes during the immune response, showed quite good interaction with each other. Next the 3D structures of the selected epitopes were generated for peptide-protein docking study. The docking was carried out to find out whether all the epitopes had the capability to bind with their respective MHC class-I and MHC class-II alleles. Since all the epitopes generated quite good docking scores, it can be concluded that, all of them had the capability to bind with their respective targets and induce potential immune response. However, among the selected epitopes, QLESKMSGK, LIRQGTDYKHWP, GVLTESNKK and TSNFRVQPTESI generated the best docking scores.

After the successful docking study, the vaccine construction was performed. The linkers were used to connect the T-cell and B-cell epitopes among themselves and also with the adjuvant sequences as well as the PADRE sequence. The vaccines, with three different adjuvants, were constructed and designated as: CV-1, CV-2 and CV-3. Since all the three vaccines were found to be antigenic, they should be able to induce good immune response. Moreover, all of them were possibly non-allergenic, they should not be able to cause any allergenic reaction within the body as per *in silico* prediction. With the highest aliphatic index of 54.97, CV-3 had the highest predicted number of aliphatic amino acids in its side chain. The highest theoretical pI of CV-1 indicated that it requires high pH to reach the isoelectric point. Quite similar results of extinction co-efficient were generated by the three vaccine constructs. The three vaccine construct showed quite good and similar results in the physicochemical property analysis.

The secondary structure of the vaccine constructs determined that CV-1 had the shortest alpha-helix formation, with 25% of the amino acids in the alpha-helix formation, however, 67.1% of the amino acids in coil formation. For this reason, most of the amino acid acids of CV-1 vaccine was predicted to be in coil structure, which was also the highest percentage of amino acids in coil structure among the three vaccines. On the other hand, CV-3 had the highest amount of amino acids in the alpha-helix formation (37.8%), according to the prediction by the study. However, all the three vaccine constructs had most of their amino acids in their coil structures. In the tertiary structure prediction, all the three vaccine construction showed quite satisfactory results. In the tertiary structure refinement and validation, CV-1 vaccine construct generated the best result with 94.3% of the amino acids in the most favored region and 4.4% of the amino acids in the additional allowed regions. CV-2 also showed good result 90.0% of the amino acids in the most favoured region. In the disulfide bond engineering experiment, only CV-1 was found to follow the selection criteria for disulfide bond formation. With the lowest and best results generated by the MM-GBSA study, HawkDock server and ClusPro 2.0 server, CV-1 was considered as the best vaccine construct among the three vaccines. For this reason, CV-1 was selected for molecular dynamics simulation study, codon adaptation and in silico coding study. The molecular dynamics simulation study was conducted by the online tool iMODS (http://imods.chaconlab.org/) revealed that the TLR-8-CV-1 docked complex should be quite stable with a good eigenvalue of 3.817339e-06. The complex had less chance of deformation and for this reason, the complex should be quite stable in the biological environment. The **Figure 10f** shows that a good number of amino acids were in the correlated motion that were marked by red color. Finally, codon adaptation and in silico cloning experiments were performed and with the predicted CAI value of 1.0, it could be concluded that the DNA sequence should have very high amount of favorable codons that should be able to express the desired amino acids in the target microorganism, *E. coli* strain K12. The DNA sequence also had quite high and good amount of GC content of 51.34%. Finally, the pET-19b vector, containing the CV-1 vaccine insert was constructed which should efficiently and effectively encode the vaccine protein in the *E. coli* cells.

The vaccine development using genome based technologies provides scientists the opportunity to develop vaccines by optimizing the target antigens. Conventional vaccines, like the attenuated vaccines or the inactivated vaccines sometimes fail to provide potential immunity towards a target antigen. Moreover, the conventional approach of vaccine development have raised many safety concerns in many pre-clinical and clinical trials. The subunit vaccines like the vaccines predicted in the study could overcome such difficulties [196]-[200]. Finally, this study recommends CV-1 as the best vaccine to be an effective worldwide treatment based on the strategies employed in the study to be triggered against SARS-CoV-2 infection. However, further *in vivo* and *in vitro* experiments are suggested to strengthen the findings of this study.

## 5. Conclusion

The SARS-CoV-2 has caused one of the deadliest outbreaks in the recent times. Prevention of the newly emerging infection is very challenging as well as mandatory. The potentiality of in silico methods can be exploited to find desired solutions with fewer trials and errors and thus saving both time and costs of the scientists. In this study, potential subunit vaccines were designed against the SARS-CoV-2 using various methods of reverse vaccinology and immunoinformatics. To design the vaccines, the highly antigenic viral proteins as well as epitopes were used. Various computational studies of the suggested vaccine constructs revealed that these vaccines might confer good immunogenic response. For this reason, if satisfactory results are achieved in various *in vivo* and *in vitro* tests and trials, these suggested vaccine constructs might be used effectively for vaccination to prevent the coronavirus infection and spreading. Therefore, our present study should help the scientists to develop potential vaccines and therapeutics against the Wuhan Novel Coronavirus 2019.

## Data Availability Statement

Authors made all the data generated during experiment and analysis available within the manuscript.

## Acknowledgements

Authors acknowledge the members of Swift Integrity Computational Lab, Dhaka, Bangladesh, a virtual platform of young researchers for their support during the preparation of the manuscript.

